# Axonemal dynein contributions to flagellar beat types and waveforms

**DOI:** 10.1101/2025.11.06.687033

**Authors:** Sophia Fochler, Matthew H. Doran, Tom Beneke, James Smith, Cecile Fort, Benjamin J. Walker, Alan Brown, Eva Gluenz, Richard J. Wheeler

**Author notes:** Correspondence to (A.B.) or (E.G.) or (R.J.W.). (T.B.): Department of Cell and Developmental Biology, Biocentre, University of Würzburg, Am Hubland, Würzburg, Germany. (J.S.): Harry Perkins Institute of Medical Research, Nedlands, WA, Australia. These authors contributed equally (Sophia Fochler, Matthew H. Doran).

## Abstract

Eukaryotic flagella, or motile cilia, are iconic molecular machines whose beating drives cell propulsion and fluid transport across diverse organisms. Beat type and waveform are tailored to function, differing between species and cell types, and individual flagella can switch between beat types. Aberrant beating causes ciliopathies and infertility in humans^1^ and prevents unicellular parasite transmission^2^. Eight distinct dynein motor protein complexes bind to axonemal doublet microtubules (DMTs) within flagella and drive beating, yet despite extensive structural analysis^3–5^, how this machinery achieves different beat types is unknown. Here, using the flagellate unicellular parasite *Leishmania*, we show a division of labour where specific dyneins drive specific beat types. Using cryo-EM, we determined the structure of the 96-nm repeat unit of the DMT and identified its dynein composition. We used CRISPR–Cas9 to systematically delete all 96-nm repeat proteins, comprehensively mapping necessity for swimming, and determined the contribution of each dynein to incidence and waveform of the preferred beat types. Outer dynein arms (ODAs) were required for symmetric tip-to-base beats, specific single-headed inner dynein arms (IDAs) were important for asymmetric base-to-tip beats (IDA*d*), and double-headed IDA*f* important for both. This systematic analysis indicates that the prevailing dogma that ODAs drive and IDAs shape the beat^6–9^ is either incomplete or not universal, and establishes new hypotheses for how different species, cell types and individual flagella achieve their necessary beat types.

## Main

Motile flagella were a key feature of the last common eukaryotic ancestor^10^ and contain a well-conserved ‘9+2’ axoneme in which nine parallel doublet microtubules (DMTs) encircle a central pair (CP) of singlet microtubules. Thousands of dyneins stud the DMTs in two rows: the outer dynein arm (ODA) and the inner dynein arm (IDA). A single ODA complex repeats every 24-nm and contains two or three heavy chains (HCs), depending on the species. IDAs are a collection of seven dyneins, each repeating every 96 nm. One contains two HCs (IDA*f*) and the remainder contain one HC (IDA*a-e* and IDA*g*). Each HC N-terminal “tail” binds auxiliary subunits and docks to the DMT, while the C-terminal “head” domain consumes ATP to drive a dynein step cycle. These dyneins drive flagellar motility by iteratively binding, pulling and releasing the adjacent DMT, resulting in sliding between neighbouring DMT pairs. Shear-resisting structures within the flagellum convert this sliding into bending.

Coordinated dynein activity establishes the beat type (tip-to-base or base-to-tip, symmetric or asymmetric) and waveform (frequency, amplitude, wavelength), all tailored to specific functions. For example, in mammals, multiciliated epithelial cell cilia undergo asymmetric base-to-tip beating to move surrounding fluid, whereas sperm use symmetric base-to-tip beating for swimming. Some flagella can alter their beat type to change function. For instance, *Leishmania* switch from symmetric tip-to-base beating for swimming to rarer asymmetric base-to-tip beating for rotation, and aberrant beating prevents progression through their lifecycle^2^.

Despite the axonemal dynein arrangement being evolutionarily conserved, diverse beating patterns occur across species. Insight into the function of individual dyneins mostly originates from *Chlamydomonas reinhardtii*, which normally undergoes asymmetric base-to-tip beating. Analysis of strains generated by random mutagenesis established the paradigm that ODAs are important for beat frequency^6^ while IDAs contribute to beat initiation and waveform shape^6–9^. However, there has been no systematic study of individual dynein contribution to achieving different beat types or waveforms. It remains unclear why such complexity of dyneins is needed and which dyneins contribute to the initiation, propagation, and modulation of flagellar beats. *Leishmania* are ideal organisms to address this, as they possess a single flagellum that switches between symmetric tip-to-base and asymmetric base-to-tip beats. Both beats are approximately planar^11^, allowing videomicroscopy and analysis as a wave in a two-dimensional focal plane^11–14^. *Leishmania* are also readily genetically modifiable, with CRISPR gene editing allowing rapid generation of deletion mutants^15,16^.

Here, we used cryo-EM to determine the structure and composition of the *Leishmania* 96 nm axonemal repeat, then systematically tested each protein for its necessity for normal cell swimming. Next, we comprehensively analysed all dyneins for their contribution to beat incidence, waveform of the tip-to-base beat, and effectiveness of the base-to-tip beat for cell rotation. This revealed that axonemal dyneins have distinct, complex functions. IDA*f* has a key regulatory function that enables flagellar beating. ODAs are important for tip-to-base beats, while also contributing asymmetry to the base-to-tip beat. Conversely, specific single-headed IDAs (particularly IDA*d*) are important for base-to-tip beats, while single-headed IDAs variously modulate wavelength, frequency and amplitude of the tip-to-base beat. Our findings therefore challenge both the simplicity and universality of prevailing dogma regarding dynein function in flagellar motility^6–9^.

### Structure-based dynein heavy chain assignment

Clade-to-clade variation in axonemal structure^3–5^ makes genus-specific structural data essential for precise functional analysis. We therefore reanalysed our cryo-EM data, which we previously used to generate an atomic model of the 48-nm DMT repeat from *Leishmania tarentolae*^17^, to reconstruct the 96-nm repeat that IDAs and regulatory complexes follow (**Extended Data Fig. 1, Extended Data Table 1, Movie S1 and SI**). The structure resolved all major axonemal dyneins (ODA and seven IDAs) (**Fig. 1a**) in pre-state conformations. Some complexes, especially radial spokes (RSs), were more poorly resolved because of on-microtubule flexibility. Using our maps and homologs of proteins independently assigned in a recent *Trypanosoma brucei* DMT structure^5^, we identified 146 DMT-associated proteins (**Table S1**).

**Fig. 1.**
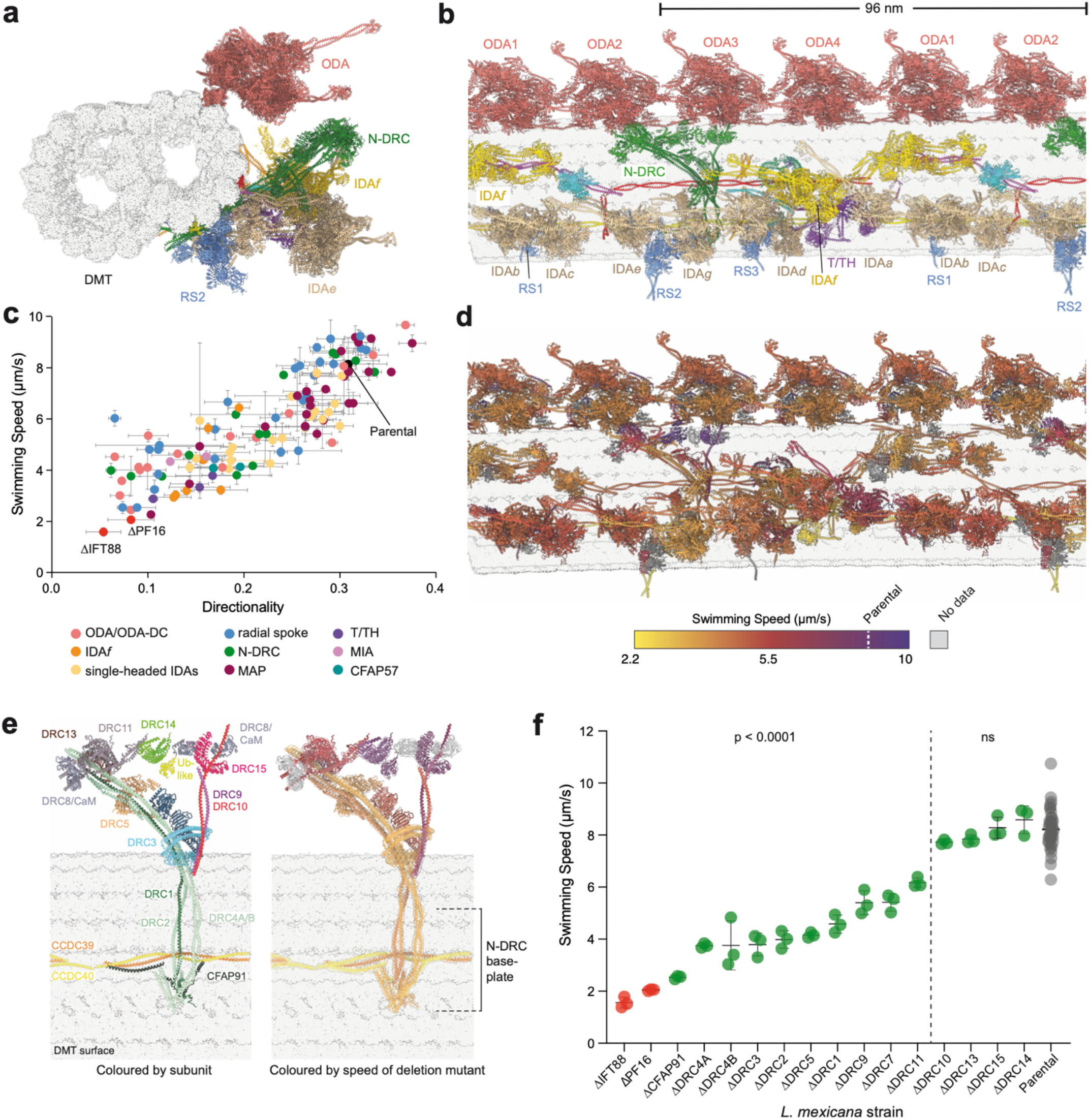
Structure-function analysis of the *Leishmania* doublet microtubule (DMT). **a-b**, Orthogonal views of the *L. tarentolae* 96 nm repeat model, with proteins colour-coded to represent key complexes: outer dynein arms (ODA, red), inner dynein arms (IDA, yellow), the nexin-dynein regulatory complex (N-DRC, green), the tether/tetherhead complex (T/TH, purple) and radial spokes (RS, blue). **c**, Quantitative analysis of swimming speed and directionality for deletions of 96 nm repeat proteins in *L. mexicana*, colour-coded as indicated in the key. Abbreviations: microtubule-associated protein (MAP, maroon), modifier of inner arms (MIA, pink). CFAP57 (teal) is an IDA*f* docking factor. Data points represent mean values, with error bars representing the standard deviation of 3 experimental replicates. ΔIFT88 and ΔPF16 are immotile controls lacking a long flagellum and with a paralysed flagellum respectively. **d**, A model of a single 96 nm repeat, with proteins colour-coded to indicate the severity of swimming speed defects associated with each deletion. Proteins in grey were not in the screen. **e**, Atomic model of the N-DRC, colour-coded by protein (left) and swimming speed of the corresponding deletion mutant (right, colour scheme as in panel d). Doublet microtubule (DMT) surface is shown in grey. CFAP91, DRC1, DRC2 and DRC4A/B form the microtubule-associated baseplate, with the binding site on the DMT dictated by the CCDC39/40 coiled coil. Swimming phenotype severity inversely correlates with distance from the microtubule: loss of distal subunits is better tolerated than loss of baseplate proteins. **f**, N-DRC deletion mutant swimming speed, ranked from most to least severe. Data points represent individual experiments; horizontal line represents the mean and error bars represent standard deviation. ΔIFT88 is an immotile control lacking a long flagellum. Significance was assessed using one-way ANOVA with Dunnett’s multiple comparisons test. *P* values: ns > 0.05, * ≤ 0.05, ** ≤ 0.01, *** ≤ 0.001, and **** ≤ 0.0001.

We identified the HCs within each dynein (**Table S2**). These localised uniformly along the length of the mature flagellum according to fluorescent or epitope tagging in the closely related species *Leishmania mexicana* (**Extended Data Fig. 2a,c-d**). Similar lack of proximal-distal asymmetry was observed for the *T. brucei* homologs using TrypTag images^18,19^ (**Extended Data Fig. 2b** and **Extended Data Fig. 3**).

The dynein distribution within the *Leishmania* 96-nm repeat is similar to that in *T. brucei* ^5^, with IDA*a* and IDA*b* associating with the base of RS1, IDA*d* and IDA*g* with RS3, and IDA*e* with DRC1 and DRC2 from the nexin-dynein regulatory complex (N-DRC). However, *Leishmania* possesses IDA*c* whereas *T. brucei* does not^5,20^, a difference explained by *T. brucei* lacking an ortholog of the *L. tarentolae* IDA*c* HC (LtaP14.1050; **Table S2**). Unlike in mammalian and *C. reinhardtii* axonemes^3,4^, IDA*c* is not RS2-associated but instead binds at the junction between CFAP189 and CCDC39/40. Notably, IDA*c* was present only in a subset of particles (**Extended Data Fig. 1e**), which originated from a subset of DMTs (**Extended Data Fig. 1f**). The absence of proximal-distal asymmetry in LmxM.14.1060 (*L. mexicana* IDA*c* HC) localisation (**Extended Data Fig. 2c,d**) suggests radial asymmetry of IDA*c* incorporation. However, our cryo-EM sample preparation loses axoneme positional information, preventing determination of the specific DMT(s) involved. We were also unable, except for LtaP25.0580 (ARM1), to identify DMT-specific densities near IDA*c*. The IDA*c*-bound region remained vacant on other DMTs (**Extended Data Fig. 1e**), leaving the mechanism underlying this asymmetry unresolved.

### Contribution of 96-nm repeat proteins to swimming

We generated deletion mutants of each 96-nm repeat protein in *L. mexicana*, expanding our previous screen of 48-nm repeat proteins^17^, and analysed their swimming speed and directionality (**Fig. 1c, Extended Data Fig. 4**, and **Table S3**). Colour-coding the atomic model by the respective deletion mutant swimming speed revealed proteins critical for motility distributed throughout the DMT (**Fig. 1d**).

The mutants exhibited a remarkable range of swimming defects, even for deletion of proteins within a single subcomplex. This is well-illustrated by the N-DRC, which is thought to be involved in electrostatic mechanosensation of its neighbouring DMT^3,21–23^. Swimming defects were more severe for deletion of DMT-bound “baseplate” proteins than for proteins located distally, toward the neighbouring DMT (**Fig. 1e, f**). Baseplate protein mutants (ΔCFAP91, ΔDRC1, ΔDRC2, ΔDRC4A and ΔDRC4B) of known importance for trypanosomatid swimming^24–27^, were among the most severe in our screen. Less severe defects in distal protein mutants (ΔDRC3, ΔDRC5, ΔDRC9, ΔDRC10 and ΔDRC11) are consistent with *C. reinhardtii* DRC7 and DRC11 mutants, where structural analysis showed maintained contact with the neighbouring DMT^23,28^.

Overall, baseplate N-DRC components are likely necessary for N-DRC assembly, while distal components may be partially functionally redundant.

### All axonemal dyneins contribute to motility

Dynein subunit deletion mutants also exhibited a wide range of swimming defects (**Fig. 1c**). Therefore, to analyse the contributions of each axonemal dynein, we focused on the HCs, deletion of which unambiguously removes motor function. To confirm that HC deletion caused selective loss of the corresponding dynein without broader axonemal disruption, we performed quantitative pairwise proteomics on axonemes from each HC deletion mutant compared to the parental strain (**Fig. 2a, Extended Data Fig. 5**, and **Table S4**). Deletion of an individual HC had little impact on the incorporation of any other HC, with two exceptions. First, ΔODA-HCα led to loss of axonemal ODA-HCβ, whereas ΔODA-HCβ retained axonemal ODA-HCα, consistent with observation that ODA-HCα is incorporated proximally in the absence of ODA-HCβ^29^. Second, deletion of either IDA*f* HC led to reduction of the other, suggesting codependence for incorporation. Thus, individual HC gene deletion results in precise loss of the targeted dynein, with the nuance that partial ODA assembly occurs in ΔODA-HCβ.

**Fig. 2.**
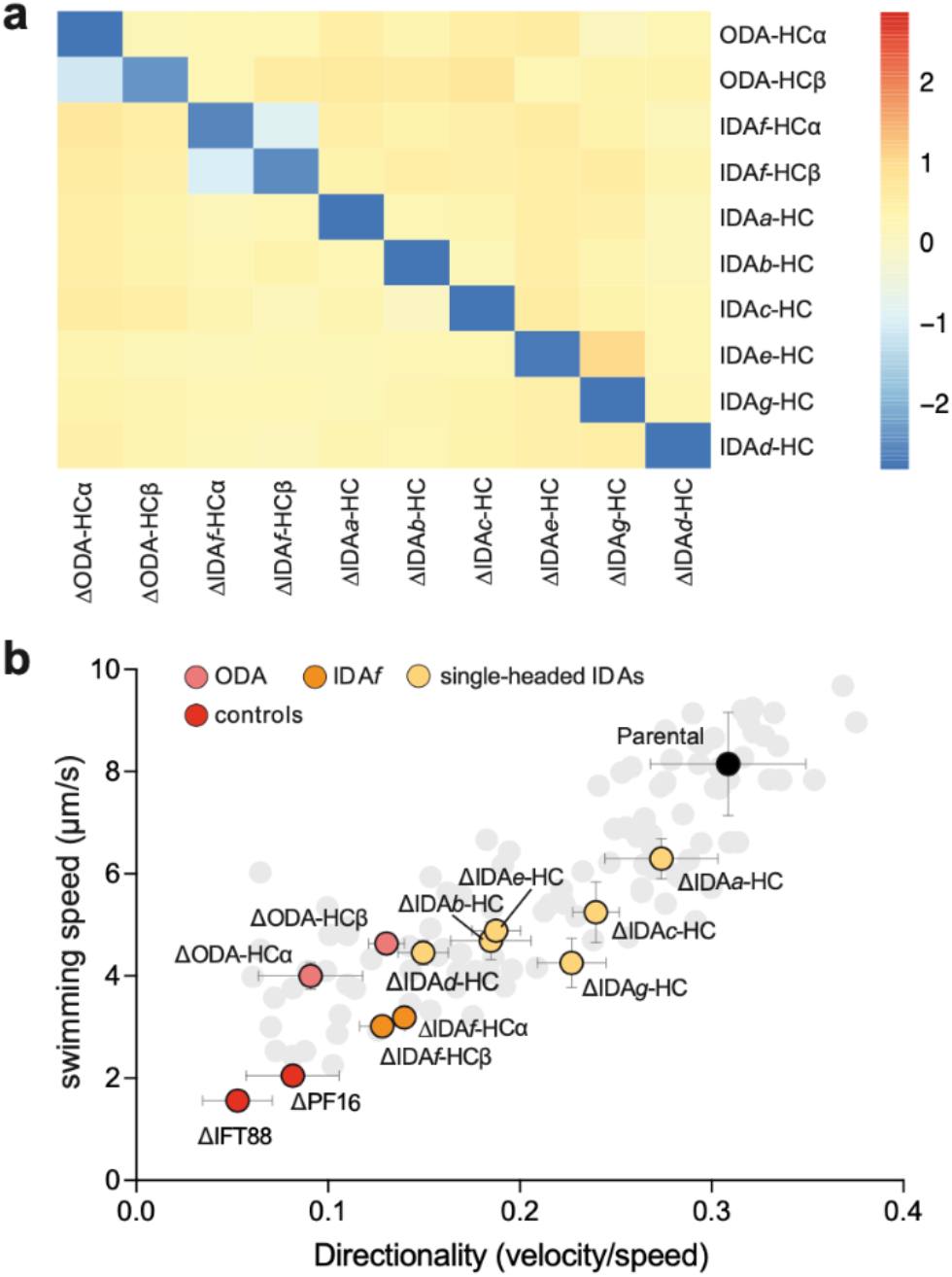
Comparative proteomic and functional analysis of axonemal dynein heavy chain (HC) deletion. **a**, Heat map summarising log_2_ fold change in flagellar HC abundance following deletion of a single HC. Except for ODA-HCα, IDA*f*-HCα, and IDA*f*-HCβ, deletion of an individual dynein HC had little to no impact on the incorporation of any other axonemal dynein HC. **b**, Dynein HC deletions highlighted within the swimming speed/directionality plot (data as in Fig. 1c). Data points represent mean values and error bars represent the standard deviation of the 3 experimental replicates. For simplicity, all other deletion mutants are shown in light grey with their error bars hidden. ΔIFT88 and ΔPF16 are immotile controls lacking a long flagellum and with a paralysed flagellum respectively.

All dynein HC deletion mutants had impaired swimming speed and directionality to varying degrees, most pronounced for ODA-HC and IDA*f*-HC deletions (**Fig. 2b, Extended Data Fig. 4b**). Contrastingly, loss of ODA-HCs particularly impacted directionality whereas loss of IDA*f*-HCs particularly impacted swimming speed. As expected, ΔIDA*f*-HCα phenocopied ΔIDA*f*-HCβ, consistent with their co-dependence, while ΔODA-HCβ swimming was less defective than ΔODA-HCα, presumably due partial ODA assembly. Among single-headed IDAs, the ΔIDA*a*-HC strain had the mildest swimming defect (small speed decrease) while other single-headed IDA mutants had comparably reduced speed and directionality. Given IDA*c* is substochiometric relative to other IDAs, the ΔIDA*c*-HC strain swimming defect was unexpectedly severe. However, it lacked a visible flagellum in ∼30% of cells (**Extended Data Fig. 6**), likely contributing to the observed severity. All other HC deletion mutants had flagellar lengths comparable to the parental strain (**Table S3**), suggesting their motility defects arise from loss of motor activity.

### Auxiliary dynein subunits necessary for normal motility

Although HC deletion did not disrupt other HCs, proteomic analysis revealed reduction of associated proteins, including docking factors, intermediate chains (ICs) and light chains (LCs), indicating HC-dependent axonemal incorporation (**Extended Data Fig. 5**). For example, in ΔIDA*f*-HC axonemes, tether/tether head complex (T/TH) subunits were reduced, consistent with known assembly dependencies in *Leishmania*^30^.

Deletion of dynein subunits may phenocopy the swimming defect of HC deletions if they are necessary for dynein assembly or normal function. We therefore analysed the corresponding deletion mutants (**Fig. 3**). For ODAs (**Fig. 3a, b**), deletion of any subunit impaired swimming, except LC4-like (**Fig. 3e**). LC4-like was the only dynein subunit whose deletion increased swimming speed, consistent with previously reported increased frequency of symmetric tip-to-base beats^31,32^.

**Fig. 3.**
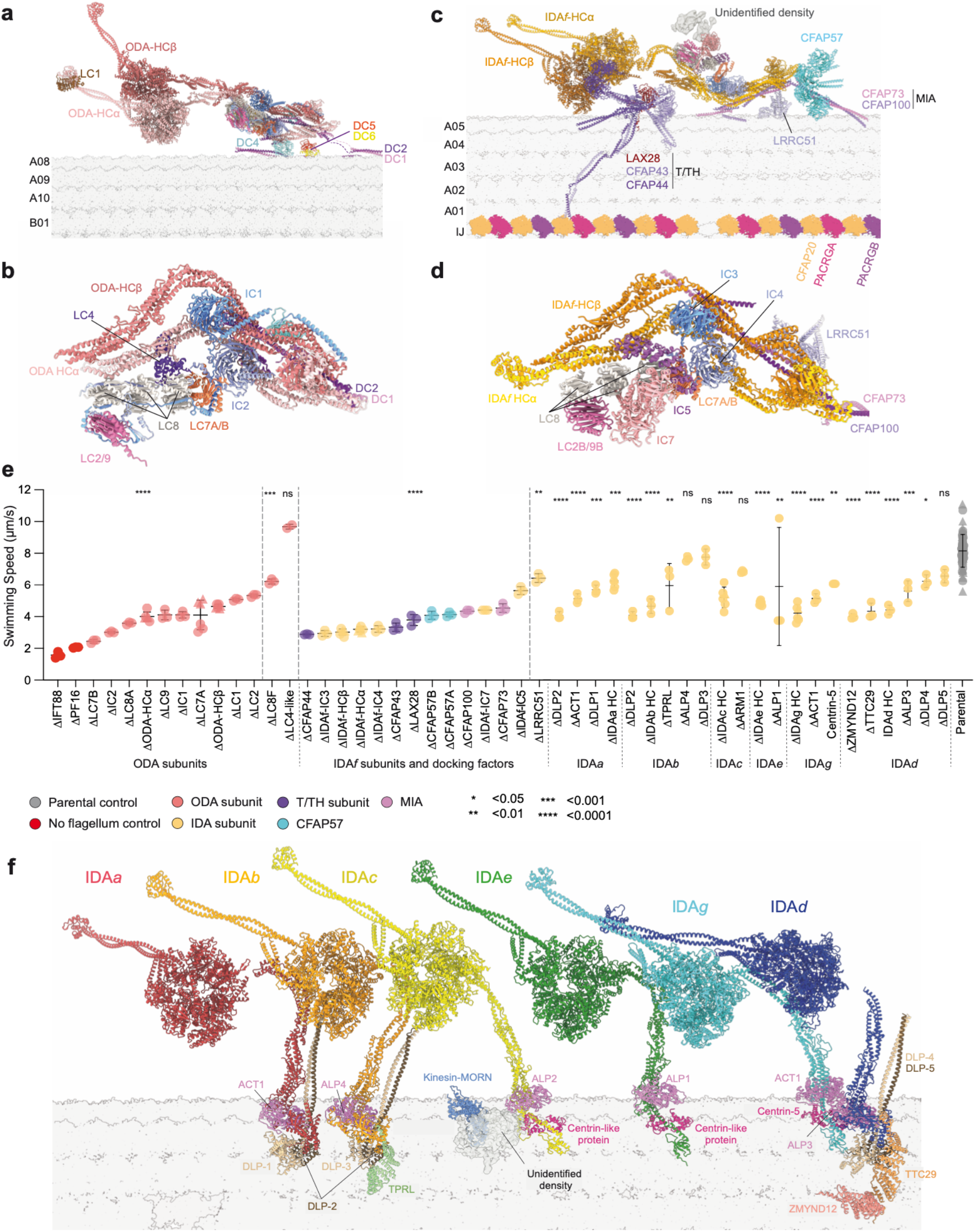
Structure and necessary features of ODAs and IDAs. **a**, Atomic model of a single ODA and its docking complex, colour-coded by protein. **b**, Subunit organisation of the ODA core. **c**, Atomic model of IDA*f* and its associated docking factors, colour-coded by protein. **d**, Subunit organisation of the IDA*f* core. **e**, Swimming speed analysis of *L. mexicana* strains lacking dynein subunits. Data points represent individual experiments, horizontal line represents the mean, and error bars represent standard deviation. Subunits ACT1 and DLP2 are found in more than one dynein complex. ΔIFT88 and ΔPF16 are immotile controls lacking a long flagellum and with a paralysed flagellum respectively. Significance was assessed using one-way ANOVA with Dunnett’s multiple comparisons test. *P* values: ns > 0.05, * ≤ 0.05, ** ≤ 0.01, *** ≤ 0.001, and **** ≤ 0.0001. **f**, Atomic models of single-headed IDAs. All other axonemal complexes have been hidden for clarity. The tail of each heavy chain is bound by an actin-like protein (ALP), and either a centrin-like protein (CLP) or a dimer of DNALI1-like proteins (DLPs). IDA*b* and IDA*d* are bound by auxiliary proteins, TPRL and TTC29–ZMYND12, respectively.

For IDA*f* (**Fig. 3c,d**), IC deletion (either IC3 or IC4), phenocopied the HC deletions, whereas other IDA*f* subunit deletions had weaker detrimental effects (**Fig. 3e**). T/TH subunit mutants (ΔCFAP43, ΔCFAP44 and ΔLAX28) broadly phenocopied the severe IDA*f*-HC deletion mutant defects (**Fig. 3e**), consistent with physical T/TH-IDA*f* association^3,33,34^ (**Fig. 3c**), linked movement during beating^4,35^, and co-dependence for axonemal incorporation^30^ (**Extended Data Fig. 5**). The T/TH may regulate IDA*f* function in addition to being essential for its assembly.

Each single-headed IDA has a unique complement of LCs including an ATP-bound actin-like protein (ALP) and either a centrin-like protein (CLP) or a dimer of DNALI1-like proteins (DLPs) (**Fig. 3f**). We identified the ALP and DLP paralog for specific IDAs based on the cryo-EM density (**SI Fig. 10**), but high centrin structural similarity restricted our identification of CLPs to IDA*g* (Centrin-5; LtaP32.0710). This single-headed IDA LC complexity is not observed for algal and mammalian species^3,4^. The resolved paralog-specific loops and terminal extensions seem to contribute to docking, suggesting that they confer specificity of IDA HC binding to its axonemal locus.

Individual DLP and ALP deletions broadly phenocopied the corresponding HC deletion (**Fig. 3e**), confirming a plausible role in docking. Deletion of proteins binding multiple HCs, for example DLP LmxM.24.0840 which binds both IDA*a* and IDA*b*, had a more severe impact consistent with disruption of both dyneins (**Fig. 3e**). Deletion of the only identified CLP (for IDA*g*) had a milder effect than deletion of IDA*g*-HC, suggesting a regulatory role (**Fig. 3e**).

In addition to established LCs, IDAs *b, c*, and *d* bind additional factors. For IDA*b* and IDA*d* these are tetratricopeptide repeat (TPR)-like proteins (TPRL and ZMYND12–TTC29, respectively) (**Fig. 3f**). Deleting these TPR-like proteins broadly phenocopied the corresponding HC deletion (**Fig. 3e**), consistent with an auxiliary docking function.

### Dynein HC contribution to preferred beat type

The observed swimming speed and directionality defects in HC deletion mutants (**Fig. 2b**) are population averages arising from multiple factors: (1) suitability of the tip-to-base beat waveform for forward swimming, (2) suitability of the base-to-tip beat waveform for cell rotation, (3) incidence of each beat type, and (4) time spent not beating. To detect changes in the incidence of specific beat types and switching behaviour among the dynein HC deletion mutants, we analysed beat incidence by manually classifying cells in 0.5-second high-speed (100 Hz), low-magnification (×20) videos. In normal culture and with 0.1% DMSO, 80% of parental cells underwent a continuous or briefly interrupted symmetric tip-to-base beat (**Fig. 4a,b**). Seven percent underwent an asymmetric base-to-tip beat, with 5% switching between waveforms during the video. The remaining cells tended to undergo slow aperiodic flagellar motion; for mutants, we also observed static curled flagella and cells lacking a visible flagellum.

**Fig. 4.**
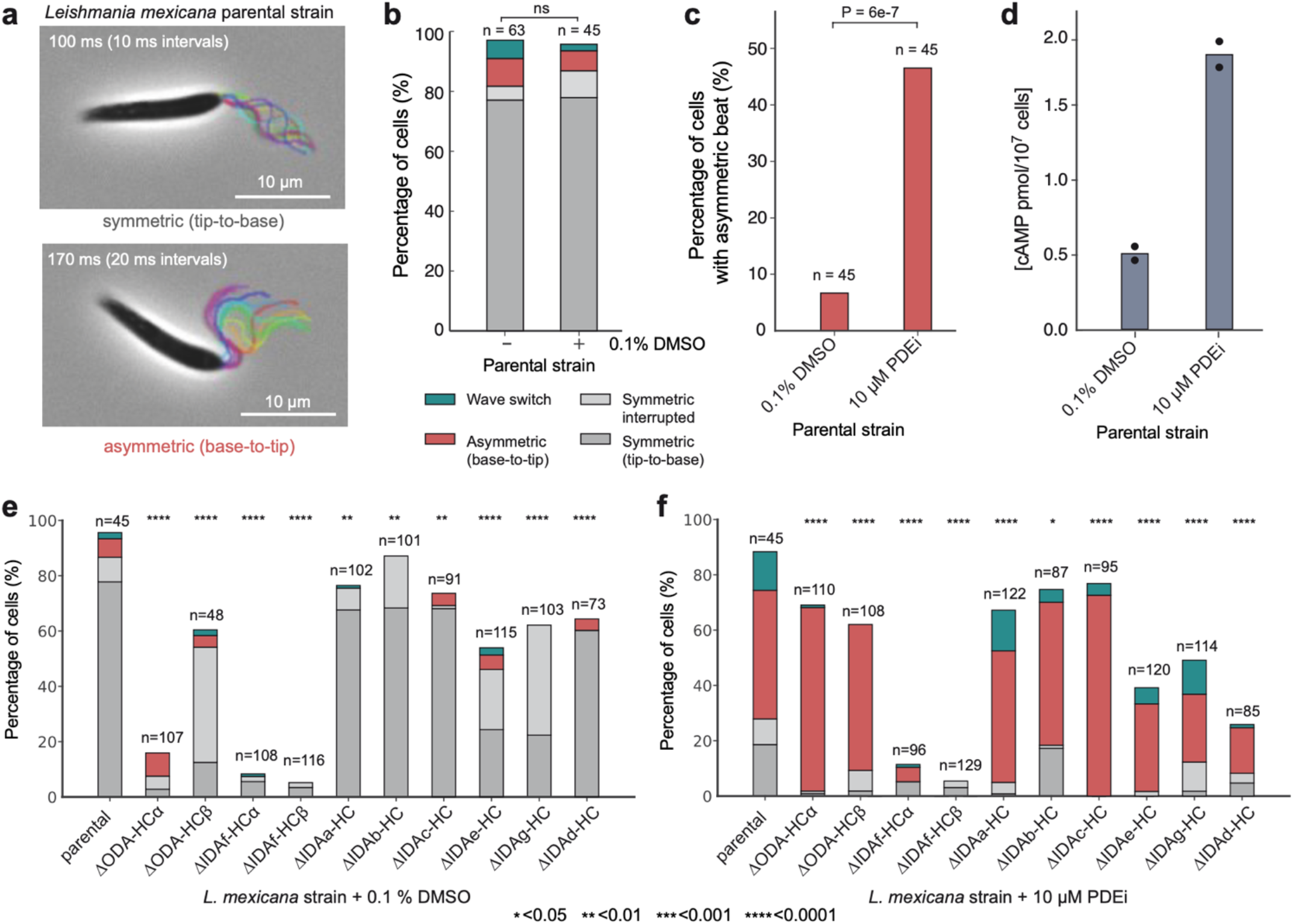
Individual contributions of dynein heavy chains (HC) to flagellar beat types in *L. mexicana*. **a**, Representative time-lapse overlays of symmetric tip-to-base and asymmetric base-to-tip beats in parental promastigotes. Movies were captured at 100 fps at 60× magnification. Flagella are rainbow colour-coded to indicate chronological progression. Scale bars, 10 μm. **b**, Beat type distribution in the parental strain population ± 0.1% DMSO. Beat types were manually classified from 0.5-s videos as undergoing: (1) continuous symmetric tip-to-base waveform (grey), (2)symmetric tip-to-base waveform with interruptions (light grey), (3) asymmetric base-to-tip waveform (red), (4) wave switch (teal). Cells without flagella or with static flagella or flagella that do not perform any recognizable beat are categorised as “other” and are not shown but account for the remaining percentage of the population. Analysis included only non-dividing cells with a single flagellum. n indicates number of cells analysed. There was no significant difference between the untreated and DMSO-treated cells (Fisher’s exact test). **c**, Proportion of parental cells with an asymmetric beat ± 10 µM phosphodiesterase inhibitor (PDEi, Compound B). *P* value was determined using Fisher’s exact test. **d**, cAMP levels increase approximately fourfold with 10 µM PDEi treatment compared to DMSO control. Bar height represents mean of technical duplicates; individual measurements shown as black circles. **e-f**, Population distributions of dynein HC mutant strains ± 0.1% DMSO (e) or 10 μM PDEi (f). The PDEi serves as a pharmacological tool to quantitatively assess the contributions of dynein HCs to the asymmetric waveform. Classes are defined as in panel b. n indicates number of cells analysed. Significance was assessed using Fisher’s exact test without multiple comparison correction. *P* values: ns > 0.05, * ≤ 0.05, ** ≤ 0.01, *** ≤ 0.001, and **** ≤ 0.0001.

Few cells naturally undergo a base-to-tip beat, which limits statistical power to detect changes in its incidence. To overcome this, we used a phosphodiesterase inhibitor (PDEi) called compound B^36^. A 10-minute incubation with 10 μM PDEi increased base-to-tip beat incidence from ∼7 to ∼47% in parental cells (**Fig. 4c**) with a fourfold increase in cellular cAMP compared to solvent control (0.1% DMSO) (**Fig. 4d**), consistent with a previous observation that cAMP induces asymmetric tip-to-base beating in demembranated *Leishmania donovani* axonemes^37^.

Specific HCs may be necessary for specific beat types or for beat switching. We tested for this by comparing parental to mutant tip-to-base beat incidence in normal culture (DMSO control) and base-to-tip beat incidence in PDEi-treated culture (**Fig. 4e,f**). No single HC deletion caused complete paralysis, therefore no individual dynein is essential for beating.

ΔODA-HCα, ΔIDA*f*-HCα and ΔIDA*f*-HCβ were almost completely unable to undergo normal symmetric tip-to-base beats. For ΔIDA*f*-HCα and ΔIDA*f*-HCβ, this was due a non-beating tightly curled proximal flagellum in many cells (76% and 58% respectively) (**Extended Data Fig. 6a,b**). The uncurled portion often underwent symmetric tip-to-base beating (**Movie S2**) but waves could not override curls to propagate into the proximal portion^38^. ΔODA-HCβ, ΔIDA*e*-HC and ΔIDA*g*-HC had more interrupted symmetric tip-to-base beats (**Fig. 4e**).

ΔIDA*f*-HCα and ΔIDA*f*-HCβ were almost completely incapable of asymmetric base-to-tip beats, and ΔIDA*e*-HC, ΔIDA*g*-HC and ΔIDA*d*-HC had reduced base-to-tip beat incidence (**Fig. 4e,f**). Interestingly, IDA*c*-HC deletion increased PDEi-induced asymmetric base-to-tip beat incidence (**Fig. 4f**), notable given that IDA*c* is likely absent from some DMTs.

Among single-headed IDAs, their roles correlated with 96-nm repeat position, with milder defects when proximal (RS1-associated IDA*a* and IDA*b*, IDA*c*) rather than distal (N-DRC-associated IDA*e*, RS3-associated IDA*d* and IDA*g*) IDAs were disrupted. ΔIDA*g*-HC and ΔIDA*e*-HC had the greatest reduction in symmetric tip-to-base beat incidence (**Fig. 4e**), whereas IDA*d*-HC had the greatest reduction of PDEi-induced asymmetric base-to-tip beat incidence (**Fig. 4f**). Overall, IDA*d-*HC deletion had the most specific effect on the asymmetric base-to-tip beat.

### Dynein HC contribution to tip-to-base waveform

Having established which HCs influence beat type incidence, we next investigated which HCs are important for specific features of the tip-to-base waveform. To do this, we automatically traced flagellar shape in high magnification (×100), high-frame rate (200 Hz) phase-contrast videos (**Extended Data Fig. 7a**).

The parental symmetric tip-to-base waveform is well modelled by sinusoidal changes in tangent angle of constant amplitude propagating from the tip at a constant speed^14^. To define this baseline, we measured amplitude, frequency, wavelength and flagellum length. Flagella have a large length range, due to growth over multiple cell cycles^39^, and strong wavelength and flagellum length correlation (R^2^ 0.89) (**Extended Data Fig. 7b**), as previously observed in related species^12^. Wavelength and flagellum length were not directly proportional, with a correlation remaining between flagellum length and residuals from a direct proportionality fit (R^2^ 0.577) (**Extended Data Fig. 7c**). Neither frequency nor amplitude correlate with flagellum length, wavelength, wavelength/flagellum length, nor to each other (**Extended Data Fig. 7c**). Thus frequency, amplitude, and wavelength/flagellum length were selected as key beat waveform descriptors with low covariance. We also calculated the time-averaged angle to measure waveform symmetry.

We measured these key descriptors in all ten dynein HC deletion strains alongside parallel parental samples, capturing flagella with tip-to-base propagating bends (**Fig. 5a,b** and **Extended Data Fig. 7a**). This was done inclusively, including low frequency less periodic motion in ΔODA-HCs and distal beating in ΔIDA*f*-HCs (**Fig. 5a**). As mutant flagella may deviate from the parental beat, we also measured goodness of fit to the single frequency and constant wave propagation speed model (**Extended Data Fig. 8a**), the constancy of wave propagation speed (phase linearity R^2^) (**Extended Data Fig. 8b**), the prominence of the dominant beat frequency (**Extended Data Fig. 8c**), and flagellum length (**Extended Data Fig. 8d**).

**Fig. 5.**
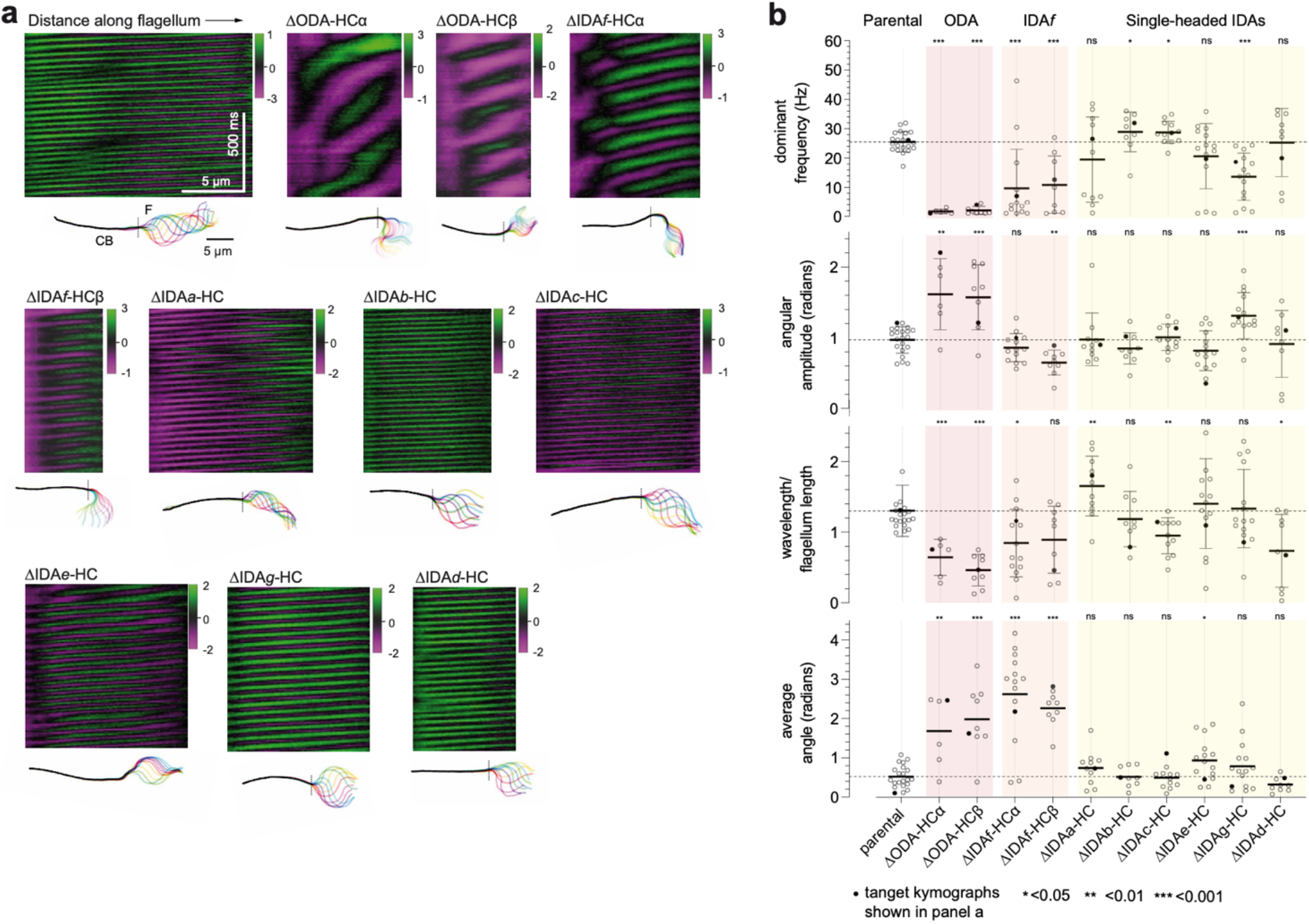
Dynein necessity for tip-to-base beat waveforms. **a**, Representative kymographs showing the tangent angle (in radians) measured at different distances along the flagellum for the parental control strain and dynein heavy chain (HC) deletion mutants. Below each kymograph are overlays of flagellar traces from a complete beat cycle, colour-coded in a rainbow spectrum to indicate chronological progression. The region to the left of the dotted line represents the cell body (CB) and the region to the right represents the flagellum (F). Scale bars, 5 μm. **b**, Waveform parameters for the dynein HC mutants. Data points in black are measurements from the kymographs shown in panel a. Statistical significance was assessed using a Mann-Whitney U test against the parental strain.

Parental beat frequency (25±3 Hz), amplitude (0.99±0.20 radians) and wavelength/flagellum length (1.34±0.40) were uniform across the population with a low average angle (0.52±0.28 radians) compared to amplitude consistent with high symmetry (**Fig. 5b**). As expected from the paradigm established in *Chlamydomonas*, ΔODA-Hcα and ΔODA-HCβ waveforms had dramatically lower frequency (**Fig. 5b, Extended Data Fig. 7d**). However, unexpectedly, the waveform also changed: amplitude increased, propagation speed became uneven, and average angle increased. Contrary to the paradigm that IDAs are necessary for a normal high beat waveform amplitude but not for a high beat frequency, the freely beating distal portion ΔIDA*f*-HC was near normal amplitude but low frequency (**Fig. 5b**). Prevalent curling or curving of the proximal region gave a high static angle.

Single-headed IDA deletion mutants had subtler waveform defects, affecting a range of parameters (**Fig. 5b, Extended Data Fig. 7f**) while still fitting the waveform model (**Extended Data Fig. 8a**). Contrasting the paradigm that IDAs define waveform shape, single-headed IDA deletion often increased (ΔIDA*b*-HC and ΔIDA*c*-HC) or decreased (ΔIDA*g*-HC) frequency (**Fig. 5b**), suggesting specific contributions to frequency regulation. However, waveform shape was also affected by single-headed IDA deletions: ΔIDA*g*-HC had increased amplitude (**Fig. 5b**), suggesting a role in amplitude regulation. ΔIDA*c*-HC had decreased and ΔIDA*a*-HC had increased wavelengths per flagellum (**Fig. 5b**), although correlation of flagellum length with wavelengths per flagellum suggests ΔIDA*c*-HC wavelength decrease is due to shorter-than-average flagella (**Extended Data Fig. 8d**), leaving ΔIDA*a*-HC as a beat wavelength regulator. Some deletions (ΔIDA*e*-HC and ΔIDA*g*-HC) also had increased asymmetry, reflected by higher average angles.

Among all deletions, ΔIDA*g*-HC had the greatest effect while ΔIDA*e*-HC and ΔIDA*d*-HC had the least effect on tip-to-base beat waveform (**Fig. 5b**). However, ΔIDA*e*-HC had reduced tip-to-base beat incidence (**Fig. 4e**), perhaps indicating a role promoting rather than shaping the tip-to-base beat.

### Dynein HC contribution to base-to-tip beats

The asymmetric base-to-tip beat rotates the cell, but the waveform is technically challenging to trace because the flagellum often crosses the cell in the focal plane. However, in the few traceable examples, we observed distinctive aberrations in HC deletions after switching from a tip-to-base beat (**Fig. 6a**). Therefore, we measured cell rotation rate after switching from a tip-to-base beat, using PDEi-treated cultures. All HC deletion mutants, except IDA*f-*HC which underwent tip-to-base beating and a subsequent switch extremely rarely, could be analysed. The productive asymmetric base-to-tip beat of the parental strain caused rotation at 45±25 degrees/s (**Fig. 6a-c**). Single-headed IDA deletions ΔIDA*a*-HC, ΔIDAe-HC, ΔIDA*g*-HC and ΔIDA*d-*HC caused reduced rotation rates (**Fig. 6c**). Most prominently, ΔIDA*d*-HC (5±4 degrees/s) (**Fig. 6a-c**) tended to switch to a paralysed state (**Movie S2**). Uniquely, ΔIDA*a*-HC flagella often continued distal tip-to-base beating while attempting proximal base-to-tip beating (**Movie S2**). Multiple IDAs therefore contribute to productive tip-to-base beats.

**Fig. 6.**
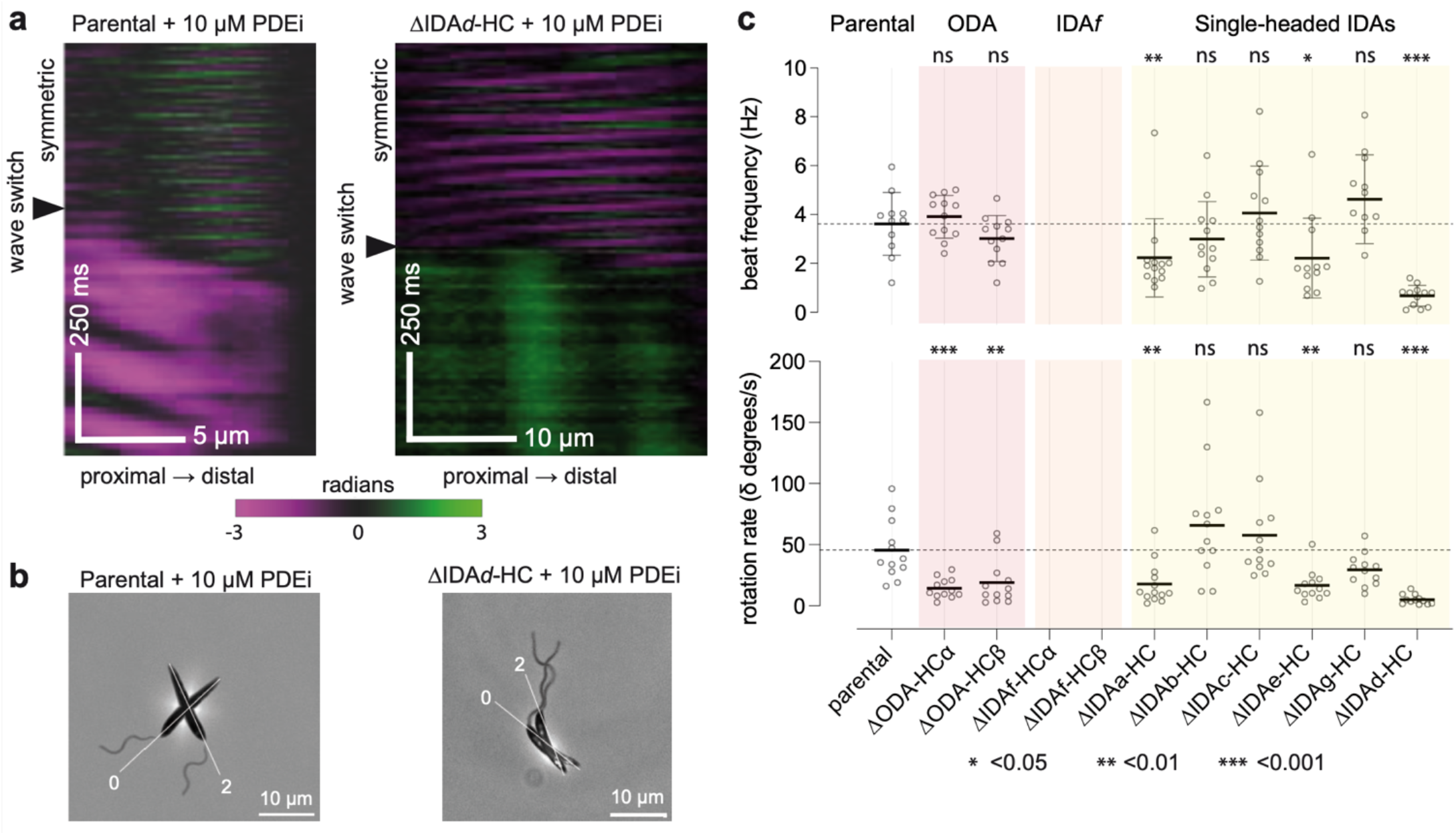
Effect of dynein HC deletions on cell reorientation. **a**, Tangent angle kymographs for a representative parental cell (left) and a ΔIDA*d*-HC mutant (right) treated with 10 μM phosphodiesterase inhibitor compound B (PDEi), capturing before and after a beat type switch from the symmetric tip-to-base beat. Scale bars, 5 and 10 μm (x axis), 500 ms (y axis). **b**, Overlay of two video frames from the cells in panel a, overlaying the time of beat type switch (0 sec) and 2 sec after. **c**, Beat frequency and rotation rate for the parental strain and each dynein HC deletion strain performing the symmetric tip-to-base beat in the presence of 10 μM PDEi. For each strain, the beat frequency (power strokes per second) and rotation rates were manually measured (n=12 cells). ΔIDA*f-*HC mutants underwent tip-to-base waveforms too rarely to analyse. Statistical significance was assessed using a Mann-Whitney U test against the parental strain.

Interestingly, ΔODA-HCα and ΔODA-HCβ also caused reduced rotation rates (16±8 and 19±19 degrees/s respectively) (**Fig. 6c**) despite high PDEi-induced base-to-tip beat incidence (**Fig. 4f**). The base-to-tip waveform instead appears more symmetric (**Movie S2**), causing slow backwards swimming rather than rotation.

## Discussion

We present the first comprehensive structure-function analysis of axonemal dyneins and their regulatory complexes, using *Leishmania* as a model organism. Through precise dynein HC deletions, we demonstrate that all axonemal dyneins contribute to swimming behaviour, with each playing distinct roles in beat type incidence, beat propagation, direction, and waveform shaping. ODAs drive symmetric tip-to-base beats but are not required for high frequency or incidence of base-to-tip beats, although they are necessary for base-to-tip waveform asymmetry. IDA*d* shows closest to the inverse function; important for base-to-tip beats but minimally contributing to tip-to-base beats. IDA*f* is a regulator of unique importance – it was the only dynein whose absence dramatically impaired *both* beats, causing severe swimming defects.

Our findings challenge the universality of the prevailing dogma, shaped by studies using *C. reinhardtii*, that IDAs define waveform shape (particularly amplitude) and ODAs drive high-frequency beating^6,40,41^. Although we cannot rule out inherent functional differences between species, the previous *Chlamydomonas* studies undersampled IDA complexity and drew conclusions from random mutagenesis that did not necessarily specifically affect HCs. In contrast, our systematic comparison used precision genetic tools and a near-complete inventory of the position and connectivity of each axonemal dynein gained from cryo-EM structures, to overcome these limitations. Furthermore, *C. reinhardtii* only undergoes base-to-tip beats, emphasising both the power of systematic analysis and considering model systems which undergo different beat types.

### Dynein roles in waveform initiation and bend propagation

Initiation of a flagellar wave requires dynein activation on one side of the axoneme, at either the tip or base, followed by coordinated propagation of this activity along the axoneme. Activation has been proposed to be spontaneous^42^ and IDA-dependent^41^. However, our data indicate that all HC deletion mutants retained some capacity to initiate both beat types (assuming that IDA*f* base-to-tip bends initiate but fail to propagate, as reasoned below), suggesting that no single dynein acts as an exclusive “waveform initiator”. However, although IDAs are sufficient to initiate waveforms, they are insufficient for normal tip-to-base initiation without ODAs, as indicated by low incidence of symmetric beats following ODA-HCα deletion (**Fig. 4e**).

We further show that while no single dynein is required for flagellum bending, IDA*f* is particularly important for bend propagation: among HC deletions, only deletion of IDA*f*-HCs greatly reduced incidence of both beat types. Furthermore, loss of IDA*f-*HCs (**Extended Data Fig. 6**), ICs or T/TH^2^ caused flagellum curling, likely due to non-IDA*f* dyneins stuck in an active state on one side of the axoneme^2^. This resembles the first basal bend during normal switching from a tip-to-base to base-to-tip beat^38^, suggesting that base-to-tip wave initiation occurs and the curled state mimics a base-to-tip wave that fails to propagate.

The reduced tip-to-base beat incidence in IDA*f* mutants may indicate importance for tip-to-base beat propagation, although it may alternatively be due to abnormally high distal DMT sliding arising from proximal curling. We overall conclude that IDA*f* (and T/TH) are important for the propagation of both beat types, certainly for the base-to-tip beat and potentially for the tip-to-base beat. This unique waveform propagation function matches its biochemical uniqueness as the only double-headed IDA and driving abnormally slow microtubule sliding *in vitro*^43,44^.

### ODAs and IDA*d* are important for specific beat propagation directions

Switching between beat types is essential for directed taxis in *Leishmania* and other organisms^45–48^, such as ODA-dependent *Chlamydomonas* photo-avoidance^40,49^. Here, we comprehensively dissected which dyneins are necessary for *Leishmania* to achieve its two beat types, revealing a division of labour. ODAs were the only dyneins necessary for a high frequency tip-to-base but dispensable for base-to-tip beating (**Fig. 5e,f**). Asymmetric base-to-tip beats are therefore predominantly driven by single-headed IDAs, for which IDA*a, e* and *g* were important, and IDA*d* was of high specific importance (**Fig. 5f**). Our data are consistent with single-headed IDAs having the inverse role of ODAs, together important for base-to-tip beating (with functional redundancy for all except IDA*d*), yet dispensable for high frequency tip-to-base beating.

Requirement of ODAs for a specific waveform is also seen in *C. reinhardtii*, where photo-avoidance uses ODA-dependent base-to-tip beats^40,49^, but ODAs are not required for asymmetric base-to-tip beats^49^ . However, without a systematic analysis of IDA function in *Chlamydomonas*, it is not possible to determine their specific contributions to symmetric base-to-tip beating.

The concept of division of labour between IDAs and ODAs is supported by divergent axonemes lacking ODAs but retaining IDAs (e.g. the eel *Anguilla rostrata* and the moss *Physcomitrella patens*)^50,51^ or vice versa (e.g. the diatom *Thalassiosira pseudonana*)^52^. These demonstrate that minimal dynein subsets can generate functional beats: IDAs or ODAs may each be sufficient to drive *a* beat, but both may be required to undergo *two* distinct beats.

Overall, these data support a model in which ODAs drive the tip-to-base beat and single-headed IDAs drive the base-to-tip beat, presumably self-defining a default frequency and waveform. Additionally, ODAs modulate the base-to-tip beat and IDAs modulate the tip-to-base beat.

### Dynein cooperation for waveform shaping

ODAs and IDAs may cooperate to enable complex, adaptable motility. We observed such cooperativity in the increased symmetry of the IDA-driven base-to-tip beat following ODA-HC deletion (**Movie S2**). While ODAs are not required for high incidence or frequency of base-to-tip beats, they are likely to be partially active as ODA deletion affected base-to-tip beat waveform, increasing symmetry. IDA*d* may mostly be inactive during tip-to-base beats, while other single-headed IDAs are likely to be partially active to shape the waveform, altering frequency and symmetry.

## Conclusion

Our integrated structural, genetic and videomicroscopy approach has enabled quantitative dissection of the individual contributions of axonemal proteins to ciliary motility. In doing so, we have generated a key resource for understanding ciliary motility and dysfunction and developed a division-of-labour model that helps explain how different waveforms are generated. The generalisability of these findings will require comparative structure-function studies in a broad range of organisms, especially those with diverse axonemal structures, beat types and waveforms. Nonetheless, we established a framework for these and other future studies to determine mechanisms of cAMP as a beat type regulator and spatiotemporal regulation of dyneins.

## Acknowledgments

Cryo-EM data of *L. tarentolae* DMTs were collected at the Harvard Cryo-EM Center for Structural Biology at Harvard Medical School (HMS). Protein mass spectrometry was performed at the University of Bern Core Facility Proteomics & Mass Spectrometry. Compound B was a kind gift of Harry de Koning (University of Glasgow). M.H.D. was funded by a BCMP-Merck Company Foundation Fellowship. T.B. was supported through Medical Research Council (MRC) PhD studentship (15/16_MSD_836338; https://mrc.ukri.org/). B. J. W. was supported by the Royal Commission for the Exhibition of 1851. A.B. was supported by NIGMS grants R01GM141109 and R01GM143183, National Science Foundation (NSF) grant 2434879, and the Smith Family Foundation.

R.J.W. was funded by a Wellcome Trust Sir Henry Dale Fellowship (211075/Z/18/Z). E.G. was supported through a Royal Society University Research Fellowship (UF160661) and grant no. MR/R000859/1 from the MRC and the UK Department for International Development (DFID) under the MRC/DFID Concordat agreement and Swiss National Science Foundation project 320030L-227939.

## Author contributions

S.F. generated and analysed *L. mexicana* deletion mutants, performed tagging experiments in *L. mexicana*, prepared axoneme samples and performed mass spectrometry analysis. M.H.D. prepared cryo-EM specimens, collected cryo-EM data, performed image analysis, and built atomic models. T.B. and J.S. generated and analysed *L. mexicana* deletion mutants. C.F. analysed *L. mexicana* deletion mutants. B.J.W. and

R.J.W. developed protocols and software to analyse flagellum beat waveforms. S.F., M.H.D., B.J.W., A.B., E.G., and R.J.W. analysed the data and wrote the paper.

## Open access statement

This research was funded in whole, or in part by the Wellcome Trust [Grant number 211075/Z/18/Z]. For the purpose of open access, the author has applied a CC BY public copyright license to any Author Accepted Manuscript version arising from this submission.

## Resource availability

### Lead contact

Further information and requests for resources should be directed to and will be fulfilled by the lead contact, Dr. Richard Wheeler.

### Materials availability

Requests for *Leishmania mexicana* deletion mutant cell lines should be directed to Prof. Eva Gluenz.

### Data and code availability

The composite cryo-EM map of the 96-nm repeat unit of DMTs from *L. tarentolae* flagella has been deposited to the Electron Microscopy Data Bank (EMDB; https://www.ebi.ac.uk/pdbe/emdb/) with accession code EMD-72629. The atomic model of the *L. tarentolae* DMT has been deposited in the Protein Data Bank (PDB; https://www.rcsb.org/) with accession code 9Y6S/pdb_00009Y6S. Each focused refinement used to generate the composite was also deposited in the EMDB with the following accession codes: EMD-71903, EMD-72432 to EMD-72454, and EMD-72563 to EMD-72622. The mass spectrometry proteomics data have been deposited to the ProteomeXchange Consortium via the PRIDE partner repository^53^ with the dataset identifier PXD069197. All custom code is available at https://github.com/zephyris/tryp-motility, https://github.com/zephyris/flagellar-beat-fourier-ij and https://github.com/mar5bar/flagella-analysis-pipeline. Any additional information required to reanalyse the data reported in this paper is available from the authors upon request.

## Materials and Methods

### Parasite culture

*Leishmania tarentolae* strain P10 (Jena Bioscience, #LT-101) promastigotes were grown in brain heart infusion (BHI) medium supplemented with 0.005% hemin chloride (Sigma, #3741) and 10 U/mL penicillin-streptomycin (Gibco, #15070063) in the dark at 26 °C following recommended procedures (Jena Bioscience).

*Leishmania mexicana* strain MNYC/BZ/62/M379 expressing Cas9 and T7 RNA polymerase^16,54^ was the parental strain for the deletion studies. Promastigotes were grown in M199 medium (Thermo Fisher Scientific, #31100027) supplemented with 2.2 g/L sodium bicarbonate at 28 °C. Cultures were maintained between 1×10^5^ and 2×10^7^ cells/mL by subculturing^15^.

### Flagella isolation

Flagella isolation from *Leishmania* cultures was performed as previously described^17^. 10 mL of exponentially growing culture was added to 200 mL BHI medium. Once at 1×10^7^ cells/mL, cells were pelleted at 1000 g for 15 min, washed with phosphate-buffered saline (PBS), and resuspended in PIPES buffer (10 mM piperazine-N,N′-bis(2-ethanesulfonic acid), 10 mM NaCl, 1 mM CaCl_2_, 1 mM MgCl_2_, 0.32 M sucrose, adjusted to pH 7.2). CaCl_2_ was added to a final concentration of 100 mM, followed by ProteaseArrest protease inhibitor cocktail (G-Bioscience, #786-108). Deflagellation was achieved by drawing the cell suspension repeatedly through a 20-gauge needle using a 10 mL syringe (Hamilton), and judged to be complete when all cells ceased swimming by light microscopy. Flagella were separated from cell bodies by centrifugation over a 33% sucrose gradient, pelleting the cell bodies. The flagella were then collected by ultracentrifugation at 100,000 g for 1 hour at 4 °C and resuspended in PMDEKP (10 mM PIPES, 25 mM KCl, 5 mM MgSO_4_, 0.5 mM EGTA, ProteaseArrest). The flagellar membrane was removed by incubating with 1% NP-40 in fresh PMDEKP, then axonemes were separated from soluble proteins and detergent by centrifugation at 10,000 g for 10 min. Finally, axonemes were splayed by incubating with 10 mM ATP-Mg^2+^ and 750 μM CaCl_2_. Cryo-EM grids were prepared with the splayed axonemes at a concentration corresponding to an absorbance at 280 nm of 6.5.

### Sample preparation for cryo-EM

Cryo-EM grids were prepared using a Vitrobot Mark IV (Thermo Fisher Scientific) at 5 °C and 100% humidity. 3 μL splayed axoneme sample was added to freshly glow discharged QUANTIFOIL holey carbon grids (R2/1, copper, 300 mesh, #Q3100CR1) and blotted with a blot force 10 for 17 s before plunging into liquid ethane.

### Cryo-EM data collection

Data for *L. tarentolae* axonemes were collected across four sessions at the Harvard Cryo-EM Center for Structural Biology. Data collection parameters are summarised in Extended Data Table 1. These datasets were reported previously and were used to determine the structure of the 48 nm repeat of the *L. tarentolae* DMT^17^. Each data collection used a Titan Krios microscope (Thermo Fisher Scientific) at 300 kV and equipped with a K3 direct electron detector (Gatan) and a BioQuantum K3 Imaging Filter (Gatan) with a slit width of 25 eV at a defocus from −0.8 to −2.5 μm. A nominal magnification of 64,000× was used, corresponding to a pixel size of 1.33 Å. 54 frame movies were collected with 5.7 s exposure time and 62.6 Å^-2^ electron dose.

### Cryo-EM image processing

Image processing was performed using RELION-5^55^ unless stated otherwise. The 96 nm repeat was resolved through 3D classification of the 48 nm repeat^17^, using a cylindrical mask covering protofilaments A02-A03-associated protein densities, yielding two well-resolved classes containing 251,436 and 281,984 particles. Each class was refined separately, using a low-pass filtered map of the 48 nm repeat as the initial reference and a box size of 512 pixels (680 Å), resulting in two halves of the 96 nm repeat at 3.7 Å resolution. Local refinement of each half, using 30 masks corresponding to 2-3 protofilaments positions on overlapping 16 nm longitudinal sections, improved the resolution to 3.0-3.6 Å. Local maps were sharpened in RELION postprocessing and with DeepEMhancer^56^. Each of these maps was then fit to their corresponding consensus map and merged into a single composite map using “vop maximum” in ChimeraX^57^. Processing schemes for resolving the microtubule-associated axonemal complexes are detailed in the **SI Methods**. Resolution estimations are based on the 0.143 FSC criterion^58^.

### Protein identification and model building

Model building was performed in Coot^59^, initiated by placing two copies of the previous 48-nm model (PDB 9E78)^17^. Proteins with 96 nm periodicity were identified and modelled using integrative methods based on map resolution. For resolutions better than 4 Å, sequences of well-built regions from ModelAngelo^60^ were extracted and searched against the *L. tarentolae* proteome using BLASTp implemented in TriTrypDB^61^. Top hits were validated against mass spectrometry analysis of the sample^17^. Confirmed hits were either extended manually in Coot or replaced with AlphaFold^62^ models. For regions with lower resolution (>4.0 Å) and known protein identities, we placed either homology models based on previous axonemal structures or AlphaFold models. In cases of unknown identity, systematic rigid body searches of AlphaFold predictions of the proteins in our mass spectrometry analysis^17^ were performed using MOLREP^59,63^. Details on the identification of specific axonemal proteins are described in the **SI Methods**.

### Model refinement

Models were refined in Coot using real-space refinement with torsion, planar peptide, and Ramachandran restraints. Trans peptide restraints were disabled to allow refinement of cis-peptide bonds where present. Because of computational limitations imposed by the size of the model, real-space-refinement in Phenix^64^ was performed on four separate submodels: (1,2) two 96 nm halves containing tubulin, MIPs and MAPs,(3) four ODA complexes, and (4) the bases of RSs and the IDA dyneins. Each submodel was refined against the composite map with secondary structure, Ramachandran, and rotamer restraints^64^. Following five macrocycles of refinement, models were manually corrected in Coot to reduce clashes and reintegrate regions that fell out of density during refinement. A final refinement with five macrocycles of global minimisation was performed. The four models were then merged and interfaces examined to fix clashes. Model validation was performed using MolProbity^65^. The final statistics are reported in Extended Data Table 1.

### Deletion mutant generation and PCR based genotyping

*L. mexicana* deletion mutants were generated and genotyped by a diagnostic PCR on genomic DNA as previously described^2^. Cell growth, flagellar assembly, and motility were qualitatively assessed by visual examination of newly generated strains and noted for comparison with subsequent quantitative analysis.

### Endogenous protein tagging

*L. mexicana* strains expressing endogenously tagged gene products were created as previously outlined^16^. For tagging with mNeonGreen (mNG), promastigotes were transfected with the DNA donor derived from the pPLOTv1 neo-mNeonGreen-neo plasmid^16^ and sgRNA constructs, then selected in M199 medium supplemented with 40 µg/mL neomycin. To tag IDAc-HC with 10TY epitopes, donor PCR products were amplified from the pPLOTv1 puro-10TY-puro plasmid^16^ and selected with 20 µg/mL puromycin. After emergence of drug-resistant populations and 2 culture passages, cells were maintained in medium without the selection drug before imaging.

### Live cell epifluorescence

Fluorescently tagged cells were prepared for live-cell imaging as previously described^29^. Exponentially growing promastigotes were washed three times by centrifugation at 800 g and resuspension in PBS. DNA was stained with 10 μg/mL Hoechst 33342 during the second wash. The washed cells were settled on glass slides and observed immediately. Widefield epifluorescence and phase-contrast micrographs were acquired using a Leica DMi8 inverted microscope with an ORCA Fusion camera (Hamamatsu Photonics) and a 63× oil objective numerical aperture (NA) 1.4.

### Immunofluorescence

Exponentially growing *L. mexicana* promastigotes (5×10^6^ cells per sample) were pelleted at 800 g and washed twice with 5 mL PBS. Washed cells were allowed to settle onto coverslips and were fixed with 2% formaldehyde (Thermo Scientific, #28906) in PBS for 10 min. After two PBS washes, cells were permeabilised with 0.1% Triton X-100 for 1 min, then washed twice with PBS. For detergent extraction of cytoskeletons, the settled cells were incubated with 0.1% Nonidet P-40 (NP-40) in PBS for 5 min followed by two PBS washes prior to fixation. Samples were blocked by incubation in PBS with 2% bovine albumin for 15 min before being incubated overnight with primary antibody BB2 (anti-Ty1 epitope)^66^ at 1:50 dilution in blocking buffer, washed thrice with PBS-Tween 20 (0.1% PBS-T), incubated with 200 µg/mL goat antimouse Alexa Fluor-488 (Thermo Fisher Scientific, #A-11001) for 2 hours, and washed thrice with PBS with 20 µg/mL Hoechst 33342 for DNA staining. Coverslips were mounted onto glass slides using glycerol. Phase-contrast and widefield epifluorescence images were acquired using a Nikon Ti2 microscope with a 100× Plan-apochromat objective (NA 1.40) and a Kinetix sCMOS camera, with identical settings for the 10TY::IDA*c*-HC and parental control samples. Images were processed using identical contrast settings in ImageJ/Fiji^67^.

### Flagellum structure distance measurements

As detailed in^17^, the distance from the kinetoplast to the onset of fluorescence in tagged *T. brucei* strains was determined from publicly available images in the TrypTag dataset^18^. Only cells with one kinetoplast and one nucleus (1K1N cells) were analysed. Strains where a large proportion of the population had very low signal intensity or where there was high cytoplasmic signal background were excluded. Controls were p197 (Tb927.10.15750) and p166 (Tb927.11.3290) as tripartite attachment complex markers^68,69^, SAS6 (Tb927.9.10550) and POC5 (Tb927.10.7600) as basal body markers^70,71^, TZP150 (Tb927.7.2150) as a transition zone marker^72^, Basalin (Tb927.7.3130) as a basal plate marker^72^, PF16 (Tb927.1.2670) and PF20 (Tb927.10.13960) as central pair markers, and PFR1 (Tb927.3.4290) and PFR2 (Tb927.8.4970) as paraflagellar rod markers.

### Cell swimming speed and directionality

Cell swimming motility assays were performed as previously described^73^. Darkfield videos (512 frame, 5 Hz framerate) of cells in 0.1-mm deep chambers were captured using a ZeissAxioimager.Z2 microscope with 10× objective and an Andor Neo 5.5 camera. Cells were automatically identified and tracks built by joining the nearest cell based on motion from the previous two frames, within a maximum distance of 15 pixels. For each tracked cell, swimming speed was calculated as *path length* divided by *path time*, and *directionality* was calculated as *displacement* (start-to-end distance) divided by path length. Means were calculated weighted by path time to avoid biases to short paths. Tracking and analysis code is available at: https://github.com/zephyris/trypmotility.

Swimming data points represent the mean of 3 independent measurements of population swimming speed and directionality, with each measurement analysing a minimum of 1000 cells per replicate. For dynein heavy chain mutants, ΔIC2 (LmxM.31.1060) and ΔODA-LC7A (LmxM.18.1010), data points represent 6 measurements, 3 from each of two separate experiments. For ΔLC4-like (LmxM.01.0620), data points represent two measurements. Quantitative swimming analysis was cross-referenced with qualitative assessments made immediately after deletion mutant generation – any mismatch was viewed as suspect and a new strain generated and data from the new replicate used. Only one strain (ΔIC2) met that criterion. Significant swimming speed differences from the parental strain were determined using one-way ANOVA with Dunnett’s multiple comparisons test.

### Beat incidence analysis

Beat incidence was determined from 1 mL of *L. mexicana* promastigotes in exponential growth (1×10^6^ to 1×10^7^ cells/mL), incubated with either 0.1% dimethyl sulfoxide (DMSO) or 10 μM phosphodiesterase inhibitor Compound B^36^ with a final concentration of 0.1% DMSO at 28 °C for 10 min. Cells were then concentrated by centrifugation at 800 g for 5 min, removal of 600–900 µL of the supernatant, then resuspension in the remaining medium. 1 µL was placed on a glass microscope slide bordered by a Dako hydrophobic pen (∼5 cm by 2 cm rectangle). A 1.0 thickness coverslip bordered by a similar hydrophobic pen rectangle was placed on top, creating a thin chamber where cells swim freely within the focal plane. 5-second phase contrast videos were captured an sCMOS Kinetix camera at 100 Hz framerate, using a 20× NA 0.3 objective on a Nikon Ti2 Inverted microscope. Beat types of cells with a single flagellum were manually categorised from the first 50 frames (0.5 s) using ImageJ/Fiji^67^ into five categories: continuous symmetric tip-to-base beat, symmetric tip-to-base beat with interruptions, asymmetric base-to-tip beat, and ‘other’ (including non-motile flagella and flagella beating with aperiodic movement).

Incidence of flagellar curling, flagellum length, and absence of visible flagella were categorised from 20× (NA 0.3) still phase contrast micrographs acquired as above, except without DMSO or PDEi. Any flagellum that crossed back on itself was categorised as curled. Flagellum length was measured manually using ImageJ/Fiji^67^.

### Base-to-tip beat analysis

Cell rotation rate and base-to-tip beat frequency were quantified from videos captured for beat incidence analysis. Twelve cells with a single flagellum undergoing a base-to-tip beat for >0.5 s were selected. To measure cell rotation, a reference line connecting the posterior and anterior poles of the cell body was marked for the start and end frames of the observed base-to-tip beating period (see Fig. 6b), and the angle between these lines was measured using ImageJ/Fiji^67^. Rotation rate was calculated by dividing the total angular displacement by observation time. Beat frequency was determined by dividing the number of beats by observation time.

### Tip-to-base beat waveform analysis

Samples were prepared as for beat incidence analysis, without DMSO or PDEi. 5-s videos were acquired with phase-contrast illumination at 200 Hz framerate using a ZeissAxioimager.Z2 microscope with 100× (1.4 NA) objective and an Andor Neo 5.5 camera. The cell and flagellum midline and width was traced automatically from a filtered and intensity thresholded image, as previously described^14^, recording coordinates for each frame having automatically identified the cell body based on width. Tracing code is available at https://github.com/zephyris/flagellar-beat-fourier-ij.

Attempted analysis of the base-to-tip waveform used the same strategy, except using PDEi-treated cells and a 40× NA 0.6 objective with the application of a 1.5× magnifier (pseudo-60× magnification).

For tip-to-base beat waveform analysis, we angle parameterised the waveform, calculating tangent angle *ϑ* at each position *l* along the flagellum arc from coordinates at each time *t* (Extended Data Fig. 7a). Wild-type symmetrical flagellar beating is well described by^74^:

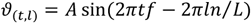

where *ϑ*_(*t,l*)_ is the tangent angle at time *t* s and position *l* μm along the flagellum arc. *A* is amplitude (in tangent angle space, radians), *f* is frequency (Hz), *L* is flagellum length (μm) and *n* is wavelength measured the flagellum arc expressed as a fraction of flagellum length (*wavelength/flagellum length*). The absolute value of *ϑ* over all positions and times is the *average angle*, which is small for the wild-type beat.

These waveform parameters were determined from the Fourier amplitude spectrum (in time) of the angle parameterised waveform, as previously reported^29^: *Frequency* was the most prominent peak and *amplitude* was aggregated over ±1 Hz of this peak. *Average angle* was calculated as the zero-frequency (static) component. *Frequency prominence* is the ratio of the amplitude contained in the most prominent peak to the total amplitude of the spectrum with the static component removed. *Phase linearity* is the R^2^ value of a linear fit of the phase of the dominant frequency as a function of position along the flagellum. A *goodness-of-fit* measure assessed the validity of single-frequency reconstruction of waveforms in Euclidian (x, y) coordinates, defined as the square root of the sum of squared error between the fitted and raw data, normalised by the sum of squares of the (x, y) coordinates of the raw data, with fitted data being rotated about the base to optimally align with raw data. Waveforms with goodness-of-fit >0.2 were rejected as unanalysable due to a poor fit. Code is available at https://github.com/mar5bar/flagella-analysis-pipeline.

For comparison of deletion mutants to the parental strain, we carried out parental beat analysis in parallel. The reference symmetric tip-to-base parental beat was defined by pooling these parental strain data with our previously published larger dataset^29^.

### Enzyme-linked immunosorbent assay (ELISA)

Exponentially growing *L. mexicana* promastigotes (3×10^7^ cells) were treated with either 0.1% DMSO (vehicle control) or 10 μM PDEi for 10 min at 28 °C. Cells were collected by centrifugation at 4 °C for 15 min at 800 g and lysed in 300 μL 0.1 M hydrochloric acid. Whole-cell cAMP concentrations were quantified using a direct cAMP ELISA Kit (Enzo Life Sciences, #ADI-900-163A), with the acetylated sample option. Technical duplicates were performed with 10^7^ cell equivalents per sample. Absorbance was measured at 405 nm using a SpectraMAX 340PC Microplate Reader (Molecular Devices). Standard curves were generated by fitting a four-parameter logistic (4PL) regression model to serial dilutions of cAMP standards using least squares fitting in GraphPad Prism. Sample cAMP concentrations were interpolated from the standard curve.

### Mass spectrometry

*L. mexicana* promastigote axonemes were isolated in triplicate from each HC mutant and parental strain. Axoneme pellets (10 μg) were resuspended in 8 M urea, reduced, alkylated and digested with sequencing grade LysC and trypsin (Promega) at 37 °C for 6 h. Digests were analysed by liquid chromatography-tandem mass spectrometry (LC-MS/MS) using a nanoElute2 UPLC coupled to a timsTOF HT mass spectrometer (Bruker). Raw data were processed with Spectronaut v.19.9 (Biognoysys) using a combined *L. mexicana* and *Bos taurus* protein database with a false discovery rate <1%. Protein quantification used a Top3 method^75^, with peptide intensities normalised by variance stabilisation^76^. Differential expression analysis was performed with appropriate statistical thresholds and imputation for missing values. See **SI Methods** for detailed sample preparation, instrument parameters, and analysis workflows.

### Data visualisation

Data were plotted using Prism v10 (GraphPad) or matplotlib.pyplot in Python v3.9.

## Extended Data: Movies

**Movie S1**. Atomic model representation of two consecutive 96-nm repeats of the *L. tarentolae* doublet microtubule. Distinct subcomplexes are colored for clarity: outer dynein arms (red), nexin-dynein regulatory complex (green), inner dynein arm *f* (yellow), single-headed inner dynein arms (straw), radial spokes (blue), and tetherhead complex (purple).

**Movie S2**. Low-magnification, high-speed videos of dynein HC deletion mutants treated with 0.1% DMSO or 10 µM PDEi.

**Extended Data Fig. 1.**
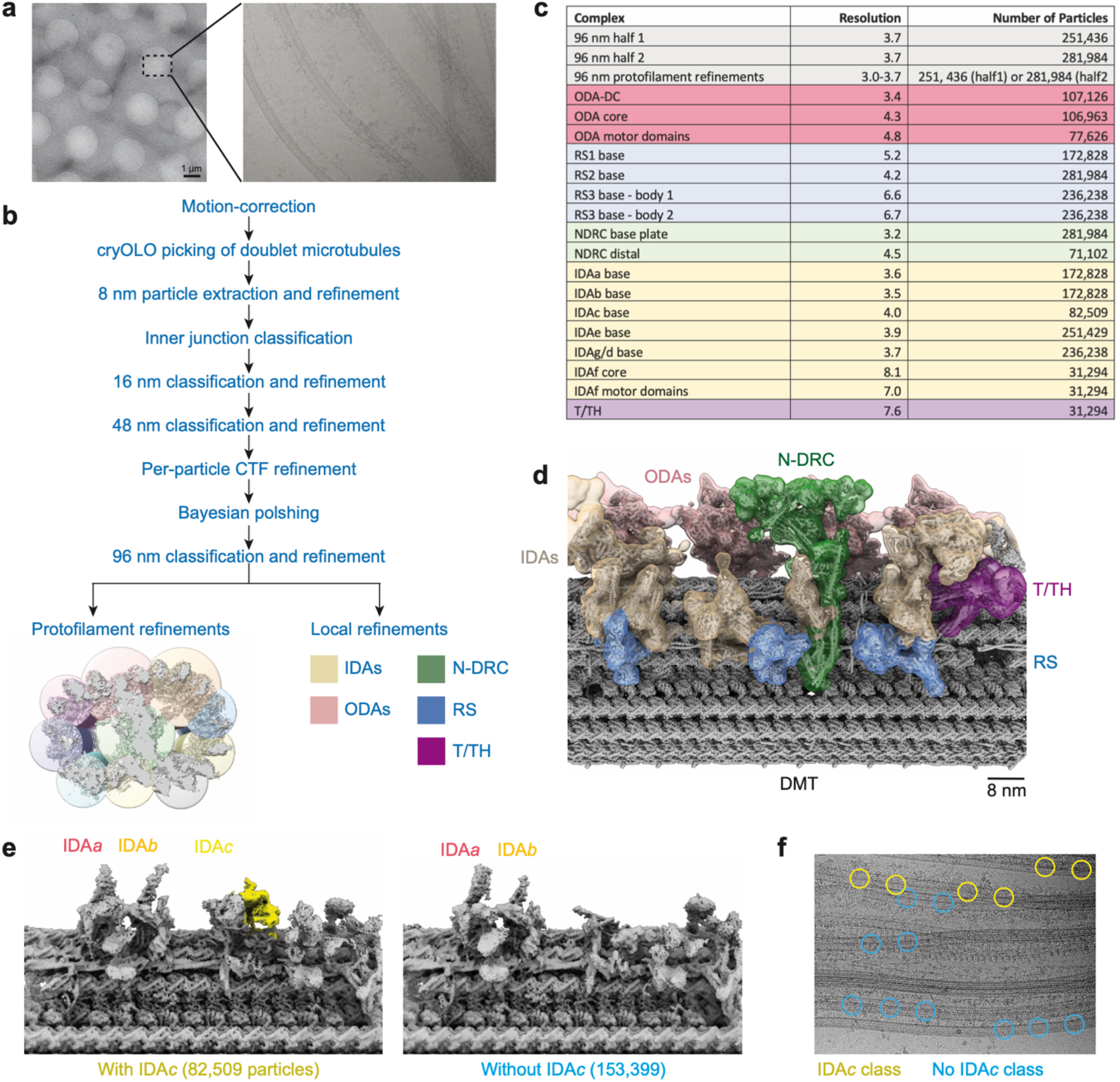
Cryo-EM processing workflow for generating the composite map of the 96-nm repeat of the *Leishmania tarentolae* doublet microtubule. **a**, Representative low-and high-magnification micrographs of splayed axonemes showing individual doublet microtubules. **b**, Simplified flow diagram outlining the key processing steps used to reconstruct a structure of the 96-nm repeat. **c**, Table summarising the resolutions of local refinements and the final particle numbers. Further details on the processing of individual axonemal complexes are provided in SI Fig. 1-9. **d**, Composite map of the 96-nm repeat with fitted atomic model. The map is coloured by axonemal complex. **e**, Cryo-EM processing identified maps corresponding to particles with and without IDA*c*. For each class, the number of particles are shown in parentheses. **f**, Visualisation of particle positions from classes with and without IDA*c* overlaid on a single micrograph. Particles segregate onto different microtubules based on their classification, suggesting IDA*c* is asymmetrically distributed on different microtubules.

**Extended Data Fig. 2.**
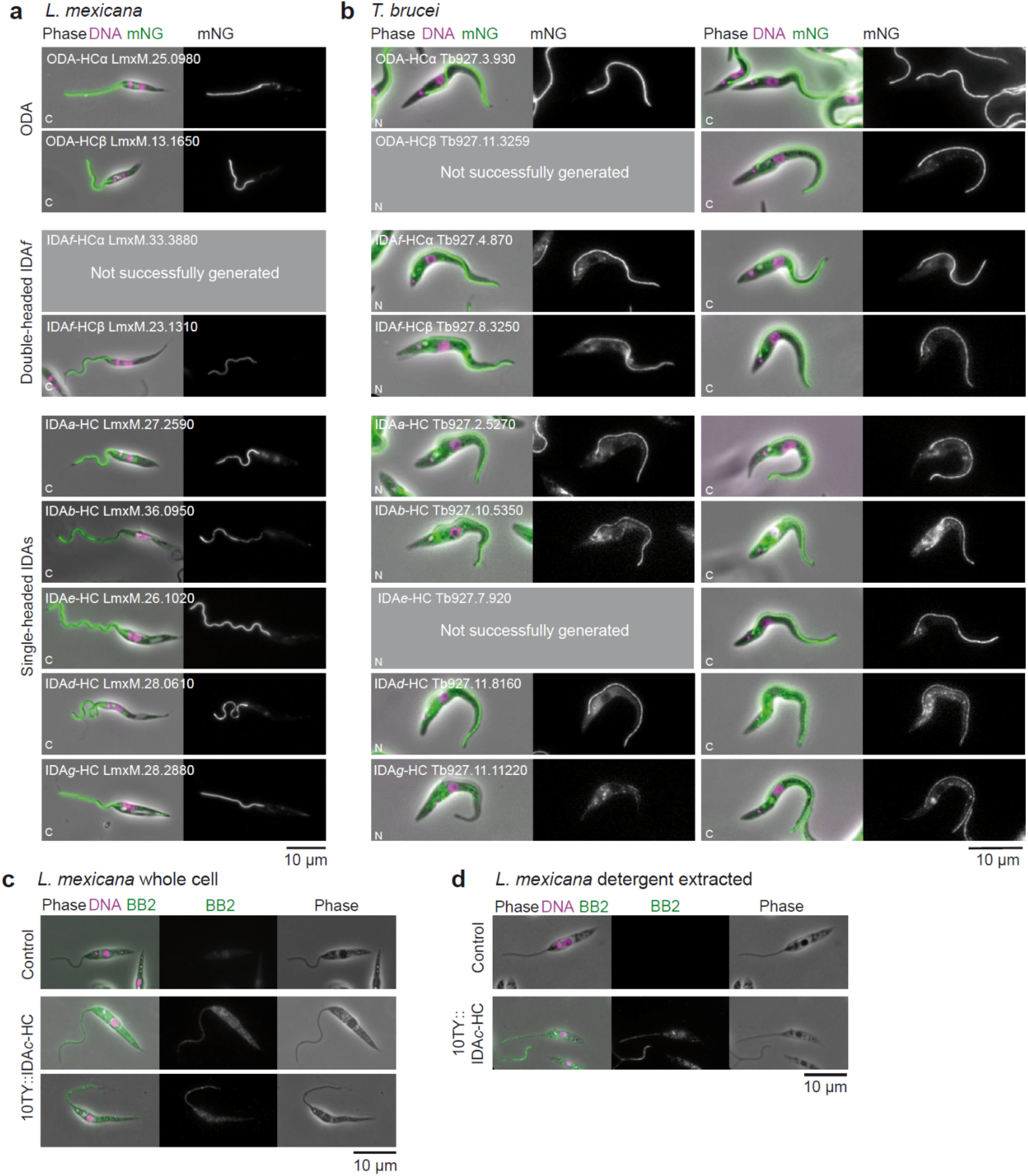
Flagellar distribution of dynein heavy chains. **a**, Localisation of dynein HCs in *L. mexicana*. Representative fluorescence images showing the cellular distribution of dynein HCs C-terminally tagged with mNeonGreen (mNG). DNA is stained with DAPI (magenta), and mNG fluorescence (green) marks the tagged HCs. Phase contrast images are shown for cellular context. Scale bar, 10 µm. **b**, Representative images of *T. brucei* cells expressing dynein HCs tagged with mNG at either their N terminus (left panels) or C terminus (right panels), sourced from TrypTag. A pair of images are shown for each tagged strain: an overlay of phase contrast (phase), the mNG fluorescence signal (green), and Hoechst 33342 DNA stain (magenta) and images that show exclusively the isolated mNG signal. Scale bar, 10 µm. **c**, Immunofluorescence localisation of IDA*c*-HC in *L. mexicana* whole cells. Cells expressing IDA*c*-HC N-terminally tagged with 10TY were immunostained with BB2 antibody (green). DNA is stained with DAPI (magenta). Phase contrast images are included. Scale bar, 10 µm. **d**, Immunofluorescence localisation of IDA*c*-HC in detergent-extracted *L. mexicana* cytoskeletons. Scale bar, 10 µm.

**Extended Data Fig. 3.**
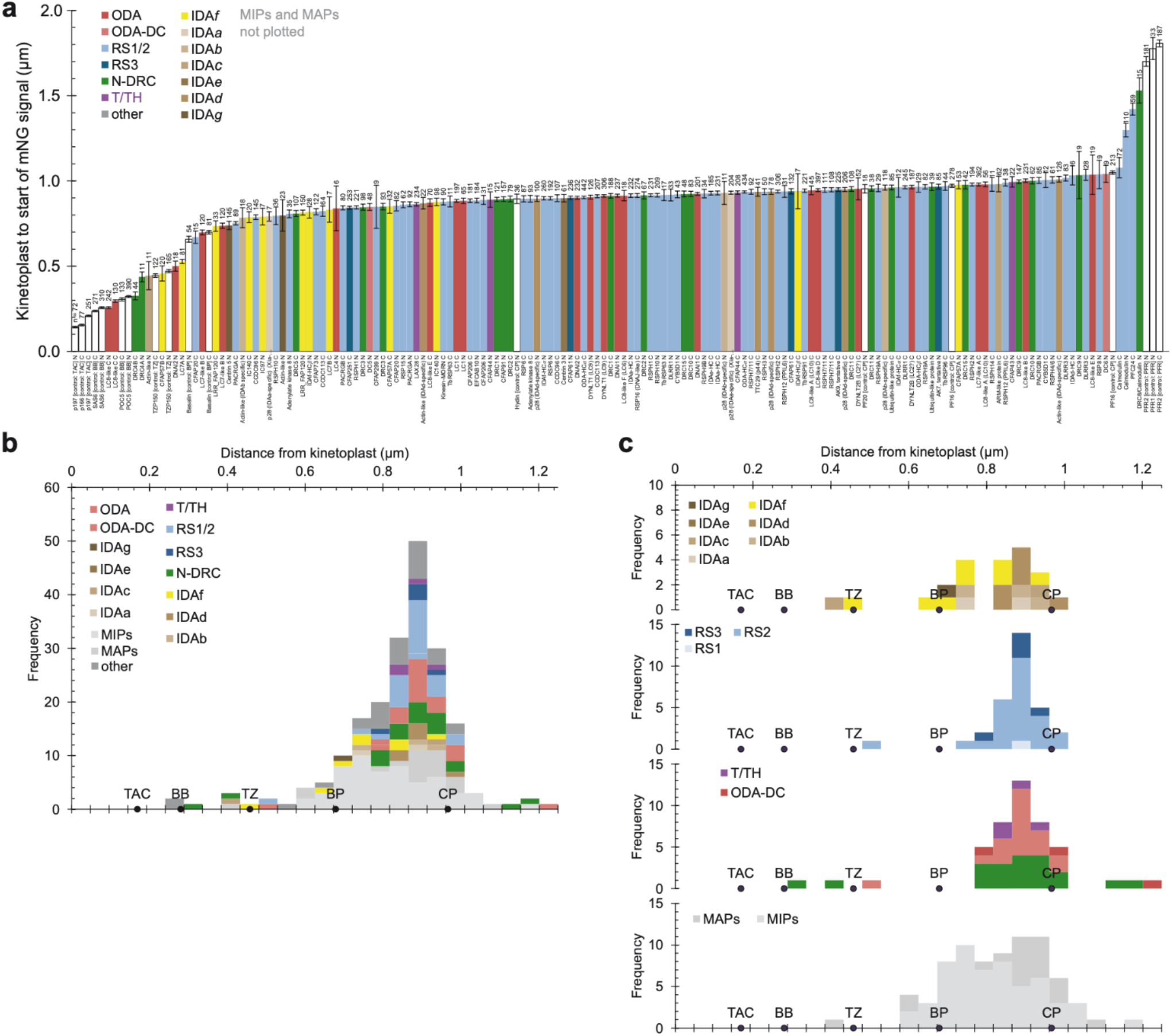
Flagellar distribution of 96 nm repeat proteins in *Trypanosoma brucei*. **a**, Automated measurement of the distance from the centroid of the kinetoplast DNA stain signal to the start of the mNeonGreen (mNG) fluorescence signal. The graph shows mean and standard error, with number of cells measured shown at the top of each bar. N and C terminally tagged strains were measured independently. Proteins are classified as indicated in the key. Controls are shown in white: p166 and p197 are components of the tripartite attachment complex (TAC); SAS6 and POC5 are components of the basal body (BB); TZP150 is a component of the transition zone; Basalin marks the basal plate (BP); Hydin, PF16 and PF20 are components of the central pair (CP); PFR1 and PFR2 are components of the paraflagellar rod (PFR). Microtubule inner proteins (MIPs) are not shown as they were plotted in^17^. **b**, A histogram summarising the localisation data shown in panel A. **c**, Histograms showing the data in panel B separated by axonemal complex type.

**Extended Data Fig. 4.**
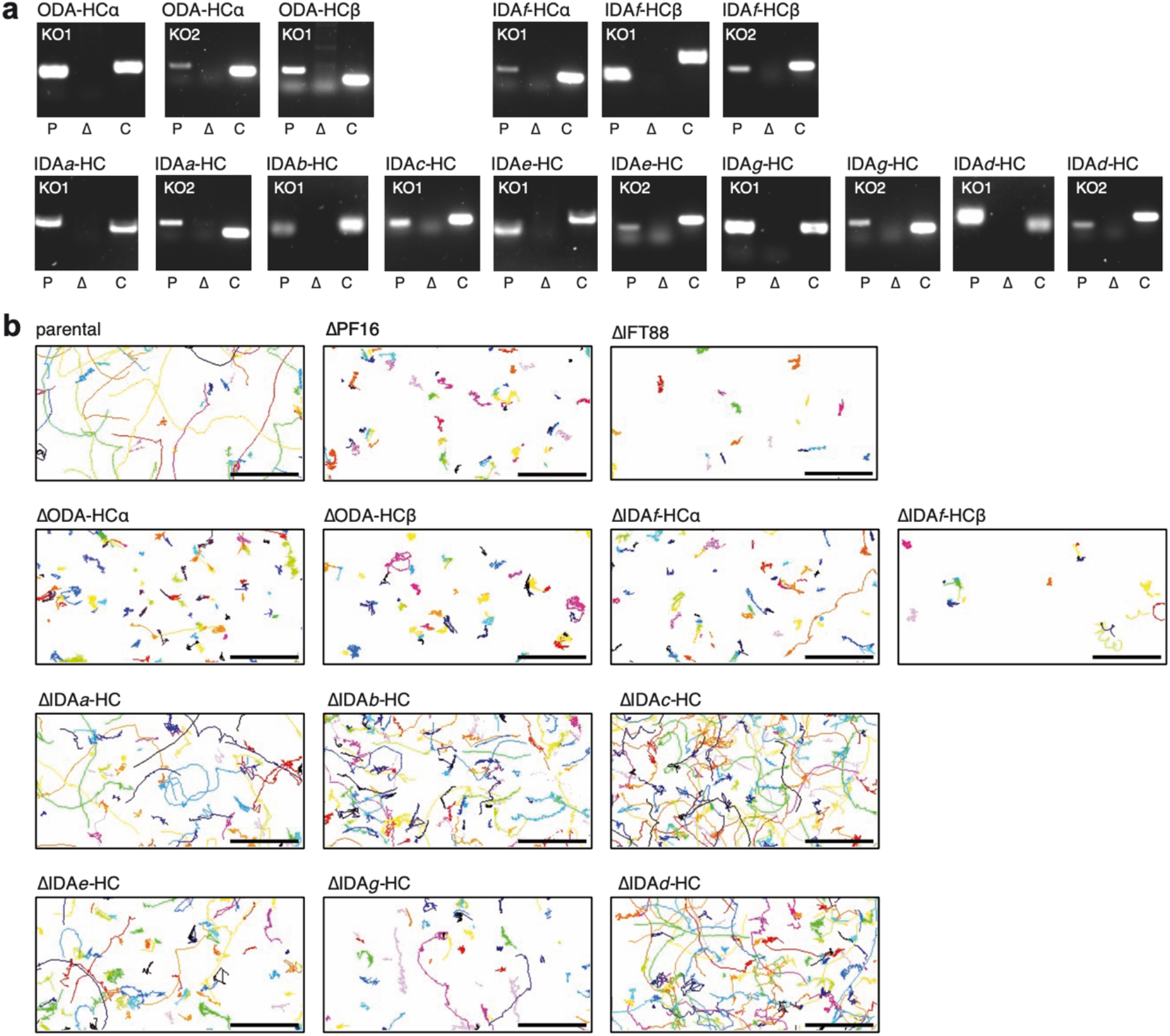
Validation and mobility of *L. mexicana* dynein heavy chain (HC) deletion strains. **a**, PCR amplification of target gene open reading frames (ORFs) visualised on agarose gels for parental *L. mexicana* Cas9 T7 cells (P) and corresponding deletion lines (Δ). A control (C) – either the ORF of PF16 or IFT88 – was amplified from the deletion strain to demonstrate the presence of genomic DNA. Absence of the target ORF PCR product was confirmed for all genes. Faint lower bands are presumed primer dimers. **b**, Swimming paths of dynein heavy chain deletion mutants extracted from darkfield microscopy timelapse videos. Each track is given a unique colour. Control strains: parental *L. mexicana* Cas9 T7 and ΔPF16 and ΔIFT88 mutants. Scale bar, 50 μm.

**Extended Data Fig. 5.**
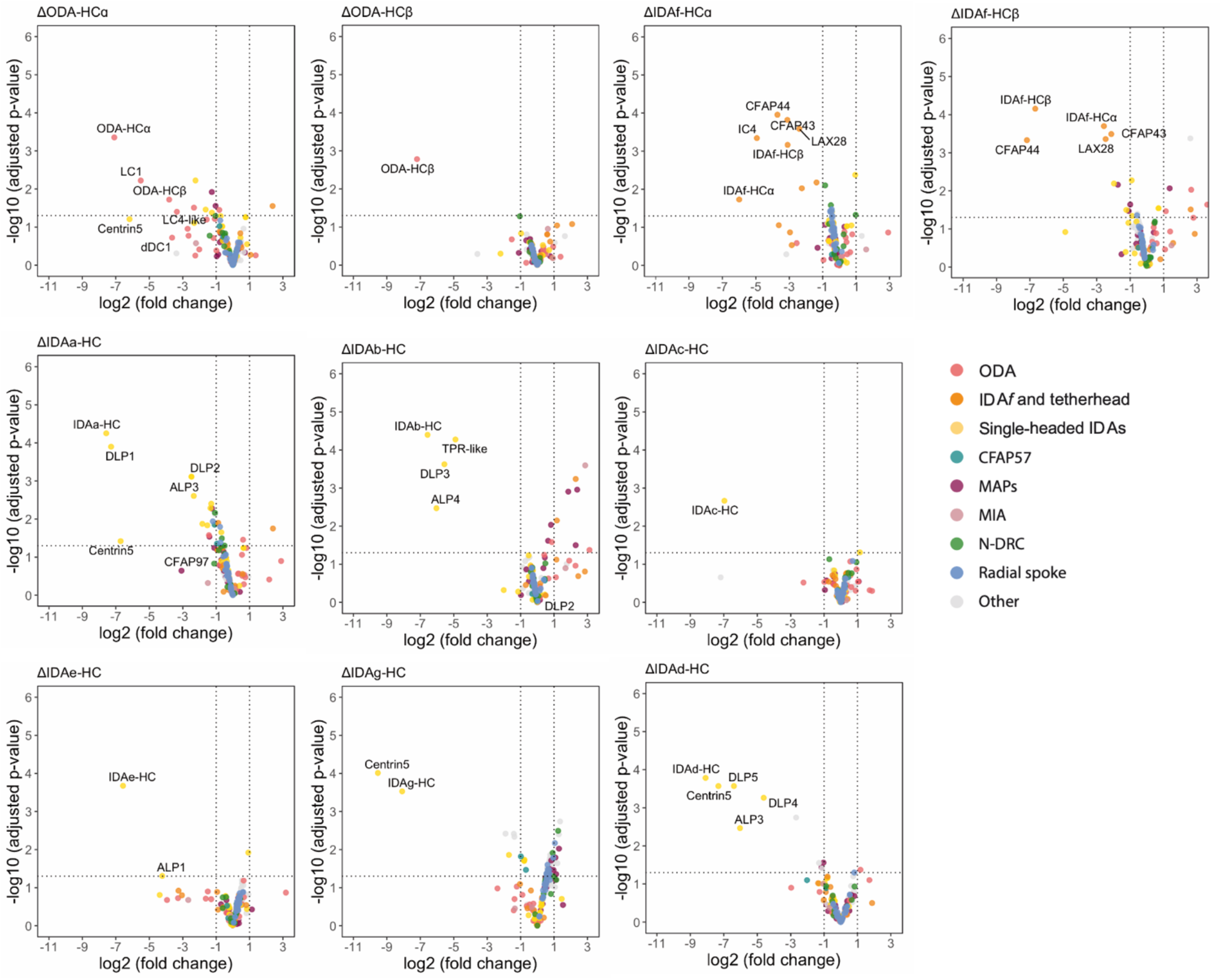
Proteomic analysis of dynein heavy chain deletion strains. Volcano plots showing differential protein abundance versus statistical significance for each axonemal dynein HC mutant compared to parental controls. Each plot represents a different mutant. Only axonemal proteins are displayed, coloured by axonemal complex as indicated in the key. Centrin-5 is decreased in multiple dynein knockout strains, suggesting it may be a binding partner of more than one dynein.

**Extended Data Fig. 6.**
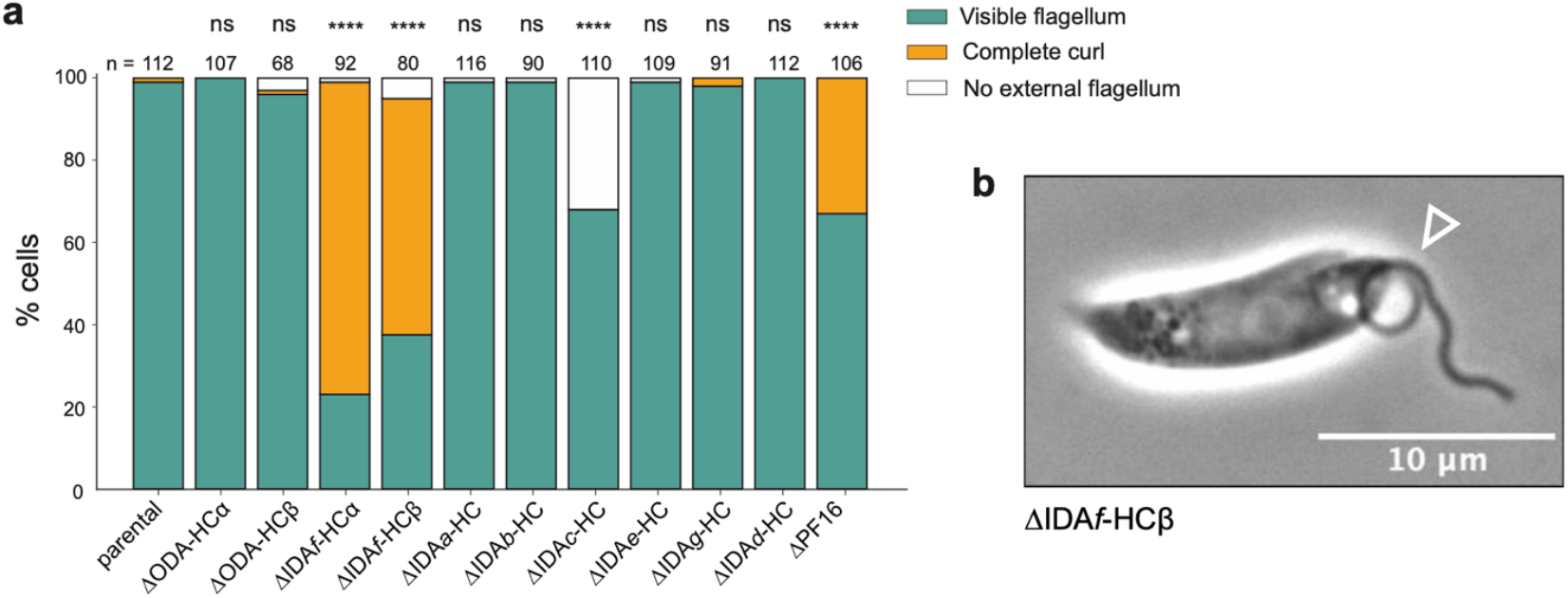
IDA*f*-HC deletion strains have curled flagella. **a**, Phenotypic assessment and quantification of curled flagella for each dynein heavy chain (HC) deletion. Controls are the parental strain and the ΔPF16 strain, which causes a central pair defect known to induce curling^2^. The number of cells analysed (n) is reported above each bar. Statistical significance was assessed using Fisher’s exact test without multiple comparison correction. *P* values: **** <0.0001; ns, not significant. **b**, Phase-contrast image of a curled flagellum in the *L. mexicana* ΔIDA*f*-HCβ mutant. The arrow marks the curled proximal part of the flagellum. Scale bar, 10 μm.

**Extended Data Fig. 7.**
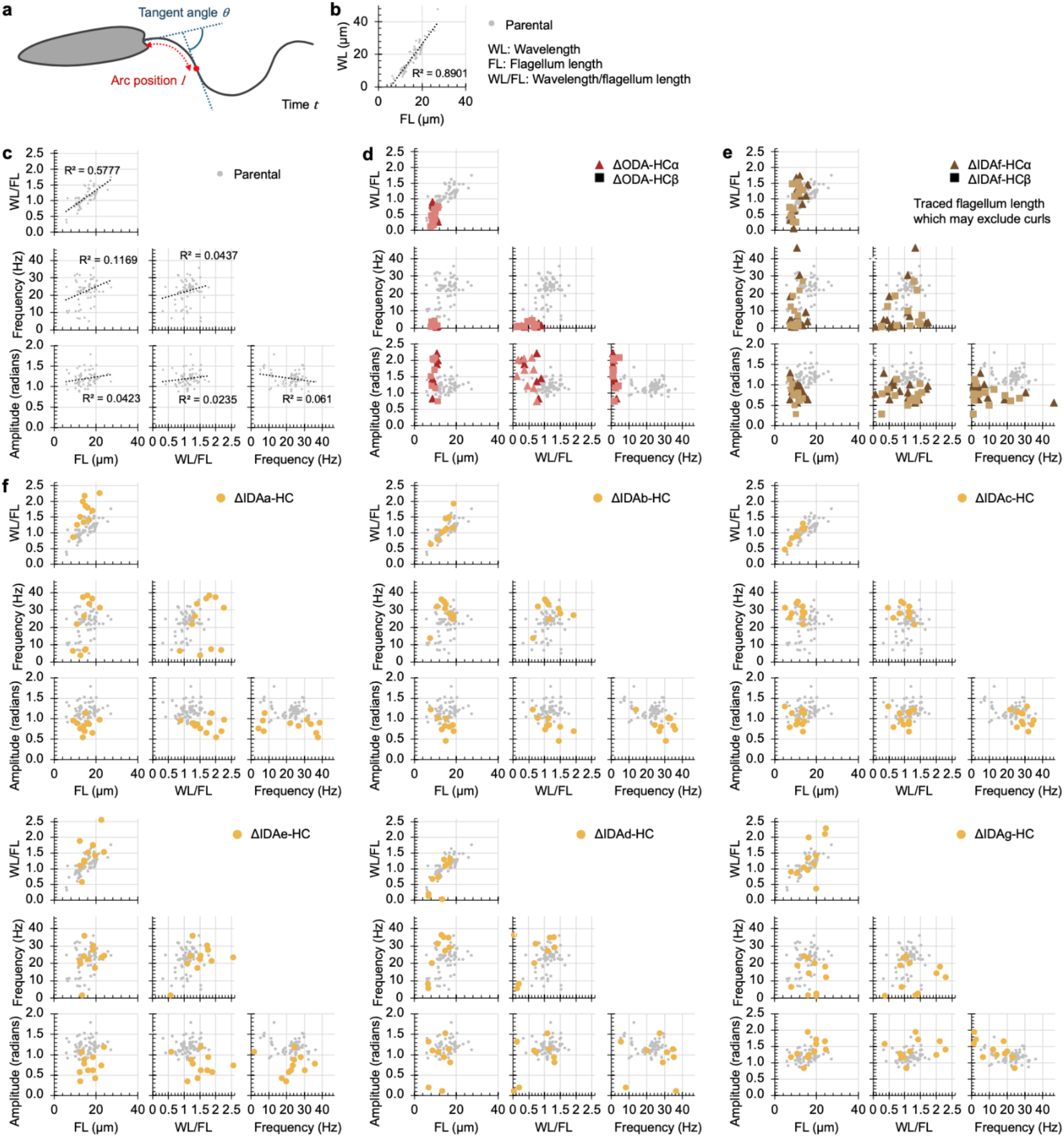
Symmetrical tip-to-base beat waveform parameters for parental and dynein HC deletion mutant beats. **a**, Diagram of the arc length and tangent angle measurements used as the basis of waveform analysis. **b**, Correlation of wavelength (WL) with flagellum length (FL) for the parental strain. Best fit line is from linear regression. **c**, Properties of the parental beat. Best fit lines are from linear regression. **d**, Beat properties of the ODA heavy chain (HC) deletion mutants. **e**, Beat properties of the IDA*f* HC deletion mutants. **f**, Beat properties of the single-headed IDA deletion mutants. Each data point represents one cell. For d-f, the parental strain data from c is plotted in the background in grey.

**Extended Data Fig. 8.**
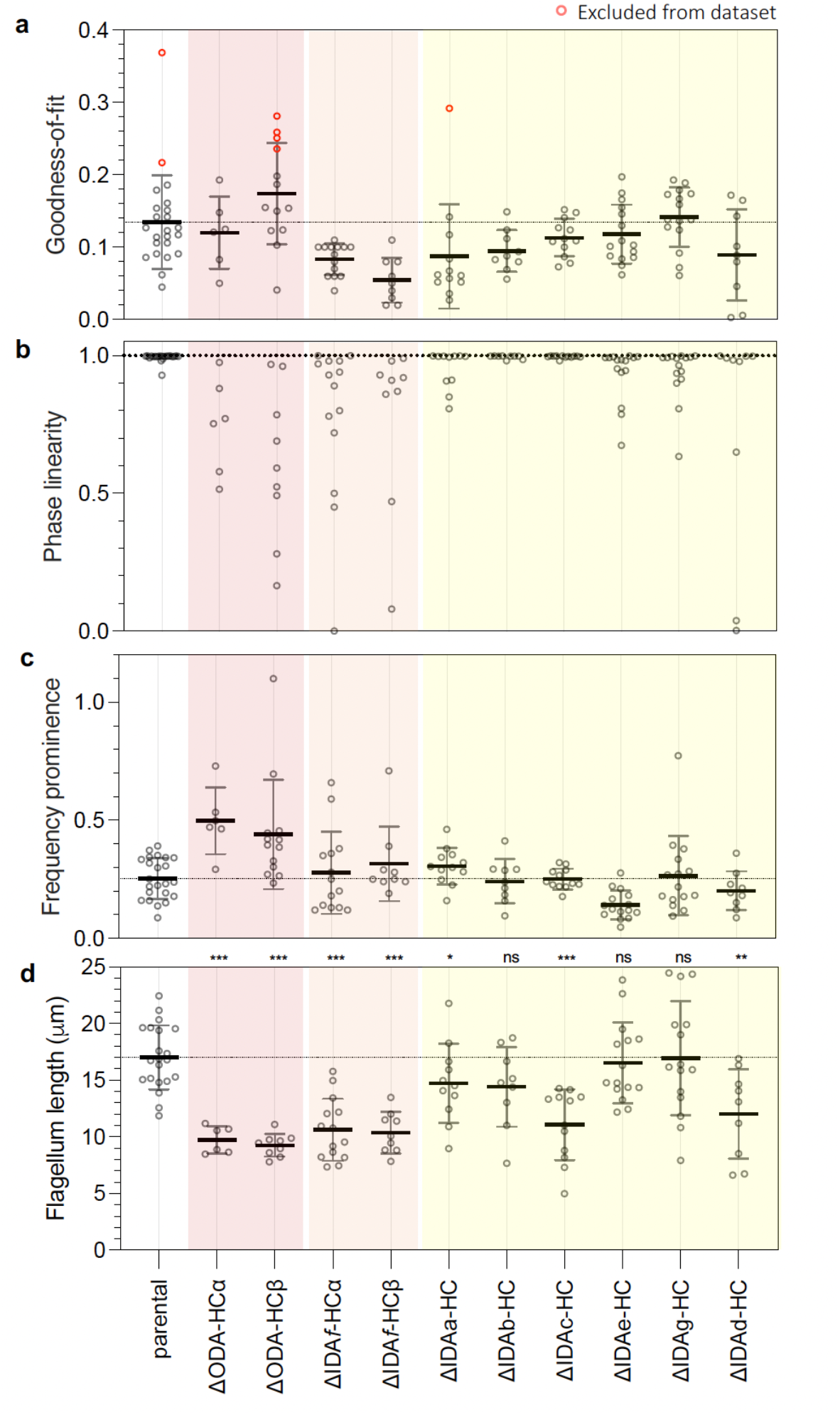
Additional metrics for tip-to-base waveforms of dynein heavy chain deletion mutants. **a**, Goodness-of-fit, where 0 is a perfect fit. A cutoff of 0.2 was manually chosen as threshold where a sufficiently good fit of the model was achieved. Red data points are out of this range, thus classified as a poor fit and these cells are excluded from other plots. **b**, Phase linearity, the quality of fit of a constant wave propagation rate down the flagellum, where 1 is constant propagation rate. **c**, Frequency prominence, proportion of frequency spectrum power in the dominant frequency peak, omitting the zero-frequency static component, where 1 represents all power being in the dominant peak. **d**, Flagellum length. Each data point represents one cell. Data points in red were excluded from the dataset due to poor goodness-of-fit, as described in Methods.

**Extended Data Table 1.** Cryo-EM data collection, refinement and validation statistics.

|  |  |
| --- | --- |
|  | *L. tarentolae* DMT, 96 nm repeat<br>(EMD-72629, PDB 9Y6S) |
| **Data collection and processing** |  |
| Facility | Harvard Cryo-EM Center for Structural Biology |
| Microscope | Titan Krios |
| Energy Filter | BioQuantum K3, 25 eV slit width (Gatan) |
| Detector | K3 (Gatan) |
| Magnification | 64,000x |
| Voltage (kV) | 300 |
| Electron exposure (e<sup>-</sup>/Å<sup>2</sup>) | 61-63 |
| Defocus range (μm) | -0.5 to -3.0 |
| Pixel size (Å) | 1.33 |
| Symmetry imposed | C1 |
| Movie stacks | 37,665 |
| Initial particle images (no.) | 7,883,514 |
| Map resolution range (Å) | 3.0-24 |
| FSC threshold | 0.143 |
| **Refinement** |  |
| Initial model used (PDB code) | 6U42 |
| Correlation coefficient (CC mask) | 0.76 |
| Map sharpening *B* factor (Å<sup>2</sup>) | 20 |
| Model composition |  |
| Non-hydrogen atoms | 3,422,763 |
| Protein residues | 432,244 |
| Ligands | 305 |
| *B* factors (Å<sup>2</sup>) |  |
| Protein | 132.92 |
| Ligand | 101.64 |
| R.m.s. deviations |  |
| Bond lengths (Å) | 0.006 |
| Bond angles (°) | 1.15 |
| Validation |  |
| MolProbity score | 1.93 |
| Clashscore | 13.35 |
| Poor rotamers (%) | 0.09 |
| Ramachandran plot |  |
| Favored (%) | 96.57 |
| Allowed (%) | 3.38 |
| Disallowed (%) | 0.05 |

\*Model validation statistics were generated by splitting the 96-nm model into three equal-sized regions and running each separately through *Comprehensive validation for cryo-EM* in Phenix. This was done because the validation job failed for the complete model due to insufficient memory.

## References

1. Wallmeier, J. et al. Motile ciliopathies. Nat Rev Dis Primers 6, 77 (2020).

2. Beneke, T. et al. Genetic dissection of a Leishmania flagellar proteome demonstrates requirement for directional motility in sand fly infections. PLoS Pathog. 15, e1007828 (2019).

3. Walton, T. et al. Axonemal structures reveal mechanoregulatory and disease mechanisms. Nature 618, 625–633 (2023).

4. Leung, M. R. et al. Structural diversity of axonemes across mammalian motile cilia. Nature 637, 1170–1177 (2025).

5. Xia, X. et al. Trypanosome doublet microtubule structures reveal flagellum assembly and motility mechanisms. Science 387, eadr3314 (2025).

6. Brokaw, C. J. & Kamiya, R. Bending patterns of Chlamydomonas flagella: IV. Mutants with defects in inner and outer dynein arms indicate differences in dynein arm function. Cell Motil. Cytoskeleton 8, 68–75 (1987).

7. Kato-Minoura, T., Hirono, M. & Kamiya, R. Chlamydomonas innerarm dynein mutant, ida5, has a mutation in an actin-encoding gene. J. Cell Biol. 137, 649–656 (1997).

8. Kubo, T., Hou, Y., Cochran, D. A., Witman, G. B. & Oda, T. A microtubule-dynein tethering complex regulates the axonemal inner dynein f (I1). Mol. Biol. Cell 29, 1060–1074 (2018).

9. Perrone, C. A., Yang, P., O’Toole, E., Sale, W. S. & Porter, M. E. The Chlamydomonas IDA7 locus encodes a 140-kDa dynein intermediate chain required to assemble the I1 inner arm complex. Mol. Biol. Cell 9, 3351–3365 (1998).

10. Richards, T. A. et al. Reconstructing the last common ancestor of all eukaryotes. PLOS Biol. 22, e3002917 (2024).

11. Walker, B. J. & Wheeler, R. J. High-speed multifocal plane fluorescence microscopy for three-dimensional visualisation of beating flagella. J. Cell Sci. 132, jcs231795 (2019).

12. Gadelha, C., Wickstead, B., McKean, P. G. & Gull, K. Basal body and flagellum mutants reveal a rotational constraint of the central pair microtubules in the axonemes of trypanosomes. J. Cell Sci. 119, 2405–2413 (2006).

13. Johnston, D. N., Silvester, N. R. & Holwill, M. E. J. An Analysis of the Shape and Propagation of Waves on the Flagellum of Crithidia Oncopelti. J. Exp. Biol. 80, 299–315 (1979).

14. Walker, B. J., Ishimoto, K. & Wheeler, R. J. Automated identification of flagella from videomicroscopy via the medial axis transform. Sci. Rep. 9, 5015 (2019).

15. Beneke, T. & Gluenz, E. LeishGEdit: A Method for Rapid Gene Knockout and Tagging Using CRISPR-Cas9. Methods Mol. Biol. (Clifton, NJ) 1971, 189–210 (2019).

16. Beneke, T. et al. A CRISPR Cas9 high-throughput genome editing toolkit for kinetoplastids. R. Soc. Open Sci. 4, 170095 (2017).

17. Doran, M. H. et al. Evolutionary adaptations of doublet microtubules in trypanosomatid parasites. Science 387, eadr5507 (2025).

18. Billington, K. et al. Genome-wide subcellular protein map for the flagellate parasite Trypanosoma brucei. Nat. Microbiol. 8, 533–547 (2023).

19. Sunter, J. D., Dean, S. & Wheeler, R. J. TrypTag.org: from images to discoveries using genome-wide protein localisation in Trypanosoma brucei. Trends Parasitol. 39, 328–331 (2023).

20. Imhof, S. et al. Cryo electron tomography with volta phase plate reveals novel structural foundations of the 96-nm axonemal repeat in the pathogen Trypanosoma brucei. elife 8, 4 (2019).

21. Bower, R. et al. The N-DRC forms a conserved biochemical complex that maintains outer doublet alignment and limits microtubule sliding in motile axonemes. Mol. Biol. Cell 24, 1134–1152 (2013).

22. Ghanaeian, A. et al. Integrated modeling of the Nexin-dynein regulatory complex reveals its regulatory mechanism. Nat. Commun. 14, 5741 (2023).

23. Kubo, T. & Oda, T. Electrostatic interaction between polyglutamylated tubulin and the nexin-dynein regulatory complex regulates flagellar motility. Mol. Biol. Cell 28, 2260–2266 (2017).

24. Hutchings, N. R., Donelson, J. E. & Hill, K. L. Trypanin is a cytoskeletal linker protein and is required for cell motility in African trypanosomes. J. Cell Biol. 156, 867–877 (2002).

25. Ralston, K. S. & Hill, K. L. Trypanin, a Component of the Flagellar Dynein Regulatory Complex, Is Essential in Bloodstream Form African Trypanosomes. PLoS Pathog. 2, e101 (2006).

26. Kabututu, Z. P., Thayer, M., Melehani, J. H. & Hill, K. L. CMF70 is a subunit of the dynein regulatory complex. J. cell Sci. 123, 3587–95 (2010).

27. Saenz-Garcia, J. L. et al. Trypanin Disruption Affects the Motility and Infectivity of the Protozoan Trypanosoma cruzi. Front. Cell. Infect. Microbiol. 11, 807236 (2022).

28. Gui, L. et al. Scaffold subunits support associated subunit assembly in the Chlamydomonas ciliary nexin-dynein regulatory complex. Proc. Natl. Acad. Sci. U.S.A. 110, 201910960 (2019).

29. Fort, C., Walker, B. J., Baert, L. & Wheeler, R. J. Proteins with proximal-distal asymmetries in axoneme localisation control flagellum beat frequency. Nat. Commun. 16, 3237 (2025).

30. Beneke, T., Banecki, K., Fochler, S. & Gluenz, E. LAX28 is required for stable assembly of the inner dynein arm f/l1 and tether/tether head complex in Leishmania flagella. J. cell Sci. 133, jcs239855 (2020).

31. Edwards, B. F. L. et al. Direction of flagellum beat propagation is controlled by proximal/distal outer dynein arm asymmetry. Proc. Natl. Acad. Sci. 115, E7341–E7350 (2018).

32. Gray, S., Fort, C. & Wheeler, R. J. Intraflagellar transport speed is sensitive to genetic and mechanical perturbations to flagellar beating. J. Cell Biol. 223, e202401154 (2024).

33. Heuser, T. et al. Cryoelectron tomography reveals doublet-specific structures and unique interactions in the I1 dynein. Proc. Natl. Acad. Sci. U.S.A. 109, E2067–76 (2012).

34. Fu, G. et al. The I1 dynein-associated tether and tether head complex is a conserved regulator of ciliary motility. Mol. Biol. Cell 29, 1048–1059 (2018).

35. Lin, J. & Nicastro, D. Asymmetric distribution and spatial switching of dynein activity generates ciliary motility. Science 360, eaar1968 (2018).

36. Gould, M. K. et al. Cyclic AMP Effectors in African Trypanosomes Revealed by Genome-Scale RNA Interference Library Screening for Resistance to the Phosphodiesterase Inhibitor CpdA. Antimicrob. Agents Chemother. 57, 4882–4893 (2013).

37. Mukhopadhyay, A. G. & Dey, C. S. Reactivation of flagellar motility in demembranated Leishmania reveals role of cAMP in flagellar wave reversal to ciliary waveform. Sci. Rep. 6, 37308 (2016).

38. Wang, Z., Beneke, T., Gluenz, E. & Wheeler, R. J. The single flagellum of Leishmania has a fixed polarisation of its asymmetric beat. J. Cell Sci. 133, jcs246637 (2020).

39. Wheeler, R. J., Gluenz, E. & Gull, K. The cell cycle of Leishmania: morphogenetic events and their implications for parasite biology. Mol. Microbiol. 79, 647–662 (2011).

40. Mitchell, D. R. & Rosenbaum, J. L. A motile Chlamydomonas flagellar mutant that lacks outer dynein arms. J. Cell Biol. 100, 1228–1234 (1985).

41. Brokaw, C. J. Control of flagellar bending: A new agenda based on dynein diversity. Cell Motil. Cytoskelet. 28, 199–204 (1994).

42. Lindemann, C. B. A model of flagellar and ciliary functioning which uses the forces transverse to the axoneme as the regulator of dynein activation. Cell Motil. Cytoskelet. 29, 141–154 (1994).

43. Kotani, N., Sakakibara, H., Burgess, S. A., Kojima, H. & Oiwa, K. Mechanical properties of inner-arm dynein-f (dynein I1) studied with in vitro motility assays. Biophys. J. 93, 886–894 (2007).

44. Kagami, O. & Kamiya, R. Translocation and rotation of microtubules caused by multiple species of Chlamydomonas inner-arm dynein. J. Cell. Sci. 103, 653–664 (1992).

45. Ringo, D. L. FLAGELLAR MOTION AND FINE STRUCTURE OF THE FLAGELLAR APPARATUS IN CHLAMYDOMONAS. J. Cell Biol. 33, 543–571 (1967).

46. Ueki, N., Matsunaga, S., Inouye, I. & Hallmann, A. How 5000 independent rowers coordinate their strokes in order to row into the sunlight: Phototaxis in the multicellular green alga Volvox. BMC Biol. 8, 103–103 (2010).

47. Shiba, K., Shibata, D. & Inaba, K. Autonomous changes in the swimming direction of sperm in the gastropod Strombus luhuanus. J. Exp. Biol. 217, 986–996 (2013).

48. Surgue, P., Hirons, M. R., Adam, J. U. & Holwill, M. E. J. Flagellar wave reversal in the kinetoplastid flagellate Crithidia oncopelti. Biol. Cell 63, 127–131 (1988).

49. Kamiya, R. & Okamoto, M. A mutant of Chlamydomonas reinhardtii that lacks the flagellar outer dynein arm but can swim. J. Cell. Sci. 74, 181–191 (1985).

50. Woolley, D. M. Studies on the eel sperm flagellum: I. The structure of the inner dynein arm complex. J. Cell Sci. 110, 85–94 (1997).

51. Hyams, J. S. & Campbell, C. J. Widespread absence of outer dynein arms in the spermatozoids of lower plants. Cell Biol. Int. Rep. 9, 841–848 (1985).

52. Idei, M. et al. Sperm ultrastructure in the diatoms Melosira and Thalassiosira and the significance of the 9 + 0 configuration. Protoplasma 250, 833–850 (2013).

53. Perez-Riverol, Y. et al. The PRIDE database at 20 years: 2025 update. Nucleic Acids Res. 53, D543–D553 (2024).

54. Beneke, T. et al. Genome sequence of Leishmania mexicana MNYC/BZ/62/M379 expressing Cas9 and T7 RNA polymerase. Wellcome Open Res. 7, 294 (2022).

55. Scheres, S. H. W. RELION: Implementation of a Bayesian approach to cryo-EM structure determination. J. Struct. Biol. 180, 519–530 (2012).

56. Sanchez-Garcia, R. et al. DeepEMhancer: a deep learning solution for cryo-EM volume post-processing. Commun. Biol. 4, 874 (2021).

57. Meng, E. C. et al. UCSF ChimeraX : Tools for structure building and analysis. Protein Sci. 32, e4792 (2023).

58. Rosenthal, P. B. & Henderson, R. Optimal determination of particle orientation, absolute hand, and contrast loss in single-particle electron cryomicroscopy. Journal of Molecular Biology 333, 721–745 (2003).

59. Brown, A. et al. Tools for macromolecular model building and refinement into electron cryo-microscopy reconstructions. Acta Crystallogr. Sect. D, Biol. Crystallogr. 71, 136–53 (2015).

60. Jamali, K. et al. Automated model building and protein identification in cryo-EM maps. Nature 628, 450–457 (2024).

61. Shanmugasundram, A. et al. TriTrypDB: An integrated functional genomics resource for kinetoplastida. PLOS Neglected Trop. Dis. 17, e0011058 (2023).

62. Jumper, J. et al. Highly accurate protein structure prediction with AlphaFold. Nature 596, 583–589 (2021).

63. Vagin, A. & Teplyakov, A. Molecular replacement with MOLREP. Acta Crystallogr. Sect. D: Biol. Crystallogr. 66, 22–25 (2010).

64. Afonine, P. V. et al. Real-space refinement in PHENIX for cryo-EM and crystallography. Acta Crystallogr. Sect. D 74, 531–544 (2018).

65. Chen, V. B. et al. MolProbity: all-atom structure validation for macromolecular crystallography. Acta Crystallogr. D Biol. Crystallogr. 66, 12–21 (2010).

66. Bastin, P., Bagherzadeh, A., Matthews, K. R. & Gull, K. A novel epitope tag system to study protein targeting and organelle biogenesis in Trypanosoma brucei. Mol. Biochem. Parasitol. 77, 235–239 (1996).

67. Schindelin, J. et al. Fiji: an open-source platform for biological-image analysis. Nat. Methods 9, 676–682 (2012).

68. Zhao, Z., Lindsay, M. E., Chowdhury, A. R., Robinson, D. R. & Englund, P. T. p166, a link between the trypanosome mitochondrial DNA and flagellum, mediates genome segregation. EMBO J. 27, 143–154 (2008).

69. Gheiratmand, L., Brasseur, A., Zhou, Q. & He, C. Y. Biochemical Characterization of the Bi-lobe Reveals a Continuous Structural Network Linking the Bi-lobe to Other Single-copied Organelles in Trypanosoma brucei *. J. Biol. Chem. 288, 3489–3499 (2013).

70. Azimzadeh, J. et al. hPOC5 is a centrin-binding protein required for assembly of full-length centrioles. J. Cell Biol. 185, 101–114 (2009).

71. Hu, H., Liu, Y., Zhou, Q., Siegel, S. & Li, Z. The Centriole Cartwheel Protein SAS-6 in Trypanosoma brucei Is Required for Probasal Body Biogenesis and Flagellum Assembly. Eukaryot. Cell 14, 898–907 (2015).

72. Dean, S., Moreira-Leite, F., Varga, V. & Gull, K. Cilium transition zone proteome reveals compartmentalization and differential dynamics of ciliopathy complexes. Proc. Natl. Acad. Sci. U.S.A. 113, E5135–43 (2016).

73. Wheeler, R. J. Use of chiral cell shape to ensure highly directional swimming in trypanosomes. PLoS Comput Biol 13, e1005353 (2017).

74. Walker, B. J., Wheeler, R. J., Ishimoto, K. & Gaffney, E. A. Boundary behaviours of Leishmania mexicana: A hydrodynamic simulation study. J. Theor. Biol. 462, 311–320 (2019).

75. Silva, J. C., Gorenstein, M. V., Li, G.-Z., Vissers, J. P. C. & Geromanos, S. J. Absolute quantification of proteins by LCMSE: a virtue of parallel MS acquisition. Mol. Cell. Proteom.: MCP 5, 144–56 (2005).

76. Huber, W., Heydebreck, A. von, Sültmann, H., Poustka, A. & Vingron, M. Variance stabilization applied to microarray data calibration and to the quantification of differential expression. Bioinformatics 18, S96–S104 (2002).

