## Supplemental Information for "Axonemal dynein contributions to flagellar beat types and waveforms"

[++]Present address (J.S.): Harry Perkins Institute of Medical Research, Nedlands, WA, Australia.

### Supplementary Methods

Cryo-EM processing of axonemal complexes

Different strategies were employed to improve the local resolution of each axonemal complex. For components near the DMT surface, including the N-DRC baseplate, IDA*f* docking factors, and the bases of the single-headed dyneins and radial spokes, refinement proceeded directly from the 96 nm halves. For components farther from the DMT surface, such as the dynein motor domains, IDA*f*, and the tetherhead complex, each 96 nm half was re-extracted with a larger box size of 660 pixels (878 Å) to accommodate the entirety of these structures. Each component was resolved individually (**SI Fig. 1-9**) using a workflow that included 3D classification to identify a subset of particles that contained the region of interest, followed by particle subtraction, recentering, and local refinement(s). These procedures were successful in resolving the bases of the single headed dyneins and radial spokes, the tetherhead complex, the motor domains of IDA*f*, as well as the N-DRC.

To obtain maps for the ODA, we merged the two expanded box-size 96 nm halves into a single particle set and performed 3D classification using a shaped mask above protofilaments A07-A08 to identify particles that contained ODA density (**SI Fig. 2**). This process identified 56,519 particles, which were refined to produce a 48 nm repeat reconstruction with prominent ODA density. To improve the ODA density further, we re-extracted particles from the 48 nm reconstruction extending +/- 24 nm and added them to the existing particle stack. This resulted in a 24 nm repeat particle stack to match the periodicity of the ODA on the DMT. Overlapping particles (6,075 duplicates) were removed to yield a final stack of 106,963 particles, which were refined to produce a reconstruction with well-resolved ODA features. The central ODA was subsequently subtracted, recentered, and the box size was reduced to 400 pixels to produce a single ODA reconstruction. A local refinement focused on the ODA core yielded a reconstruction at 4.2 Å resolution. To resolve the motor domains, an additional classification was done focused on the region above the core. This yielded one class with suitable density containing 77,626 particles. A final refinement improved the resolution of the motor domain to 4.8 Å.

To obtain maps for IDA*c*, we applied a shape mask over diffuse density present between RS1 and RS2 in the 96-nm maps and performed 3D classification without alignment (**SI Fig. 5**). A single class (out of 6) with prominent density was selected and used for refinement, generating a map consistent with a single-headed dynein HC. To improve the density adjacent to IDA*c*, the IDA*c* density was subtracted, centered, and the box size shrunk. Subsequent refinements improved the resolution to 4.0 Å.

Protein identification and model building

ModelAngelo[1] was used to identify most microtubule-associated proteins, N-DRC baseplate proteins, as well as the heavy chains, actin-like proteins (ALPs), and DNALI1-like proteins (DLPs) of IDA*a,* IDA*b*, IDA*d*, and IDA*g*. Examples of this process are provided in **SI Fig. 10**. ModelAngelo also identified Centrin-5 (LtaP32.0710) at the base of IDA*g*. Centrin-like proteins (CLPs) are also identified associated with IDA*c* and IDA*e*, but the maps are not sufficiently resolved to allow unambiguous assignment. These have been tentatively modeled as LtaP07.0720 and LtaP36.6270 in the deposited PDB file (PDB:9Y6S).

For the N-DRC, regions nearest the microtubule surface were unambiguously assigned using ModelAngelo to trace sequences through the density. For regions farther from the tubulin surface, we used homology modeling based on the *C. reinhardtii* N-DRC (PDB:8GLV)[2]. The remaining density was mostly made up of density resembling leucine rich repeat (LRR) proteins. To assign these densities, we created a focused AlphaFold library of LRR proteins identified by mass spectrometry. Each protein within the library (approximately 15) was docked as a rigid body to each of the LRR densities using MOLREP[3] . For each density, a clear top hit was established whose size and shape matched with the selected density. We named these proteins by ascending molecular weight as DRC13, DRC14, DRC15 (LtaP27.1370, LtaP09.1550, and LtaP05.0160 respectively). Residual density within the N-DRC was identified as consistent with a ubiquitin-like protein using a MOLREP search against the BALBES database of protein domains[4,5]. The top BLAST hit for ubiquitin in the *L. tarentolae* proteome, LtaP30.2640, was subsequently fitted into this region, however, in the absence of sidechain density, the assignment remains tentative.

<u>Dynein identification and modelling</u>

The ODA was modeled using homologs of subunits identified through mass spectrometry analysis of purified *T. brucei* ODA[6]. Initial models for the ODA-HCs were generated by homology modeling based on the bovine sperm ODA (PDB:9FQR)[7]. Initial models for the intermediate and light chains (except LC1 and LC4) were generated by predicting a single complex using AlphaFold3[8]. LC1 and LC4 were predicted separately.

The IDA*f*-HCs were modeled using homology models with the bovine sperm IDA*f* (PDB:9FQR)[7] acting as a template. IDA*f* intermediate chains IC3 (also known as DNAI3 or IC140), IC4 (DNAI4 or IC138), IC7 (DNAI7 or IC97), and FAP120 were identified by BLASTp searches from the bovine sperm homologs. Unambiguous identification of the light chains was not possible. We therefore modeled LC7A/B as chains LtaP32.3750 and LtaP31.3640 respectively as both are LC7 homologs not assigned to the ODA[6]. Following the same rationale, we assigned the Tctex-type light chains as LtaP35.2930 and LtaP05.0430. LC8 paralogs could not be distinguished, so we built 3 dimers of LC8C (LtaP32.0240) into our model. To create a starting model for model building, sequences for the intermediate and light chains were input into AlphaFold3 and individually rigid body fit into our density.

The heavy chains, ALPs, and DLPs of IDA*a*, IDA*b*, IDA*d*, and IDA*g* were identified from well-resolved regions of density using ModelAngelo. The IDA*e* HC was identified as LtaP26.9050 based on fit to the density and an AlphaFold3 prediction that it could interact with the N-terminal residues of DRC1/2. The IDA*e* ALP (LtaP15.1240) was identified as the only ALP with an N-terminal extension that matched with the density associated with IDA*e*. The IDA*c* HC was identified as LtaP14.1050 by a process of elimination and the lack of a corresponding homolog in *T. brucei*, which lacks IDA*c*. LtaP25.0580 was identified as part of the density adjacent to *IDAc* using a MOLREP search with our AlphaFold library. Further density in this region could not be identified.

<u>Mass spectrometry</u>

*L. mexicana* promastigote axonemes were isolated in triplicate from a minimum of $2x10^8$ cells per strain as previously reported[9]. Protein concentration was determined using a bicinchoninic acid (BCA) protein assay kit (Pierce, #A65453). Pellets containing 10 µg of protein were resuspended in 12 µL of 100 mM Tris-HCl (pH 8.0), 8 M urea. Protein concentration was reassessed with the Qubit assay (Thermo Fisher Scientific). Samples were reduced, alkylated and digested with sequencing grade LysC and trypsin (Promega) as described[10] with the exception that LysC at a protein-to-protease ratio of 100:1 and trypsin at a 50:1 ratio were added together, and digestion was carried out at 37 °C for 6 hours. For each digest, a 2 µL aliquot ($1/26^{th}$ of about 8 µg protein digest) were analysed on a LC-MS/MS system consisting of a nanoElute2 UPLC and a timsTOF HT mass spectrometer (Bruker) with chromatography and acquisition parameters as described[11].

Raw mass spectrometry data were analysed with Spectronaut v. 19.9 (Biognosys). The protein search database combined sequences from *L. mexicana*, *Bos taurus*, and common contaminants. The hybrid directDIA+ (Deep) mode was applied with default settings, except that precursor qvalue, precursor PEP, protein qvalue and protein PEP cutoffs were set to 0.01. Single-peptide protein identifications were discarded. Carbamidomethyl (C) was set as a fixed modification. Variable modifications included N-terminal acetylation and methionine oxidation. Trypsin was specified as the digestion enzyme with a maximum of 2 missed cleavages allowed.

Proteins flagged as contaminants were excluded from further analysis. The distribution of fragment peak areas reported by Spectronaut was inspected in R using the *iq* package (v.1.10.1)[12]. Fragments with the lowest intensities or not used for quantification were removed. Protein quantification was performed using the Top3 method[13]: peptide intensities were calculated from the sum of fragment intensities and normalized by variance stabilization[14]. For each protein, the same set of top three peptides were chosen for all samples based on the sum of intensity across samples. Proteins with fewer than 2 quantified peptides in a sample were classified as not detected.

Differential protein abundance was assessed using moderated t-tests implemented in the R package *limma*, with multiple testing correction performed using the Benjamini-Hochberg method. Proteins were included only if they were met minimum detection thresholds: at least 2 quantified replicates per group for experiments with three replicates per condition, or at least 5 quantified replicates per group for experiments with nine replicates per condition.

Missing values were imputed using a two-step approach at the protein group level for Spectronaut and IQ measures, and at peptide level for Top3 measures. If there was at most one non-zero value in the replicate group for a protein group, then the missing values were imputed by drawing random values from a Gaussian distribution of width 0.3 × sample standard deviation centred at the sample distribution mean minus 2.5 × sample standard deviation at protein level, respectively 2.8 × sample standard deviation at peptide level. Remaining missing values were imputed using maximum likelihood estimation[15]. Results were validated across 20 independent imputation cycles to ensure robustness. Statistical significance was determined using an adaptive p-value threshold that became more stringent for smaller absolute $\log_2$ fold changes.

### Supplementary Figures 1 to 10 for Supplementary Methods

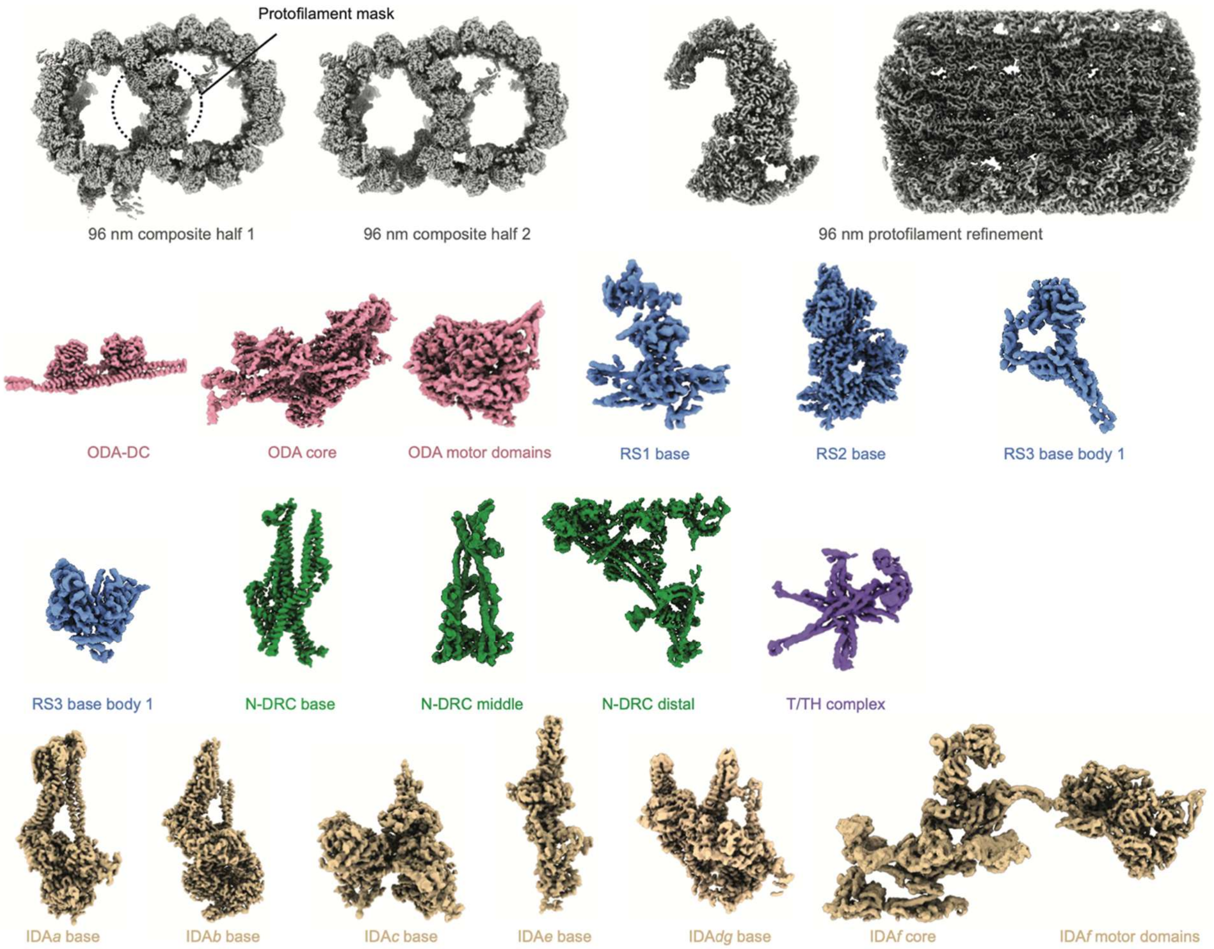

**Supplementary Figure 1 | Overview of local resolutions.**
Local masked refinements were applied to improve the quality and resolution of individual 96 nm regions of the axoneme. The final map is shown for each local refinement. These locally refined maps were subsequently docked into global consensus maps to generate the final composite 96 nm reconstruction.

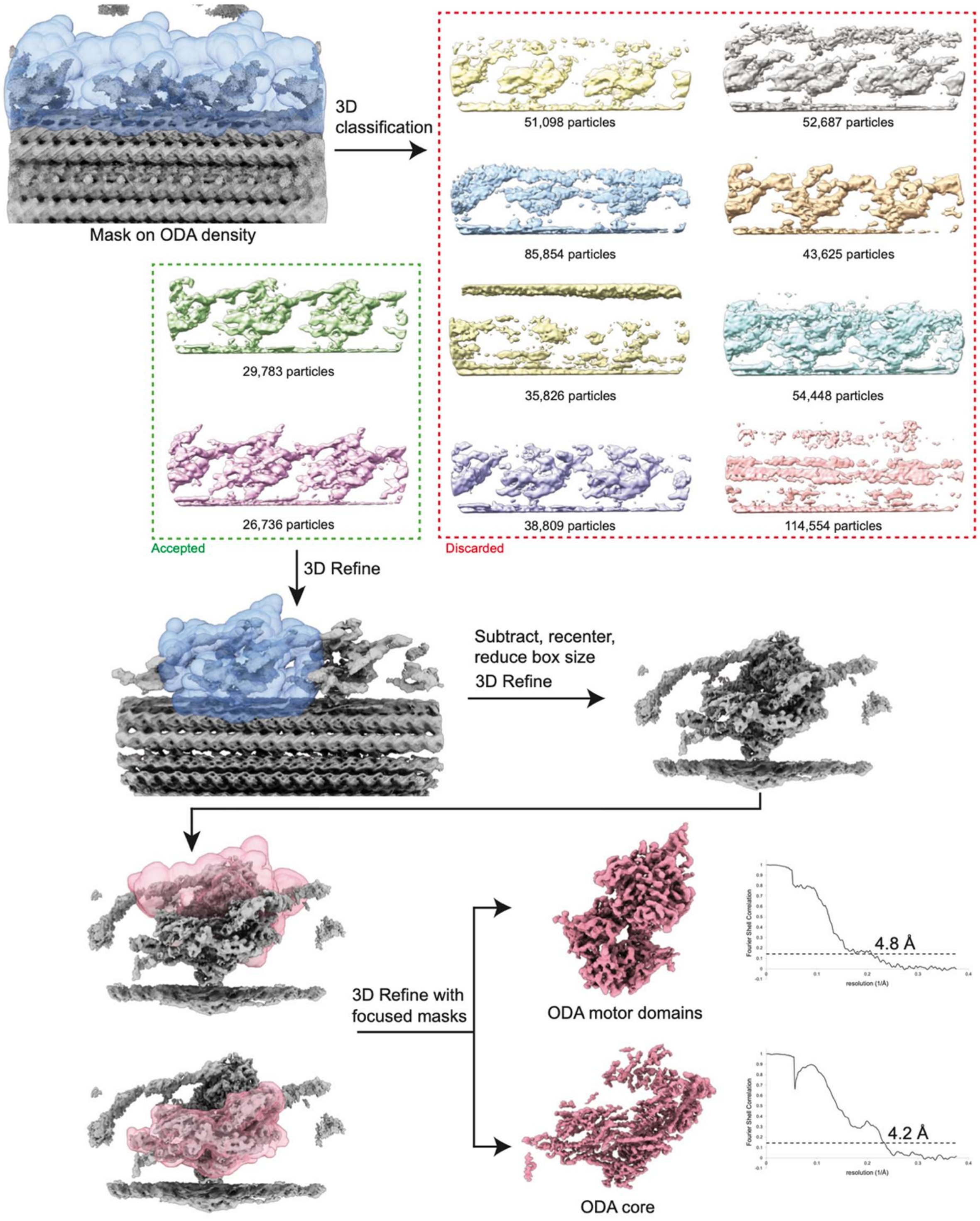

**Supplementary Figure 2 | Processing strategy for outer dynein arms (ODA).**
A total of 533,420 “48-nm particles” were subjected to 3D classification using a mask applied above protofilaments A07–A08 of the doublet microtubule (DMT). Classification yielded two well-resolved classes containing ODA-bound particles that were combined and refined. Particles for each ODA were recentered and re-refined. Mask-focused 3D local refinements were then used to improve the maps of the motor domains and the ODA core region.

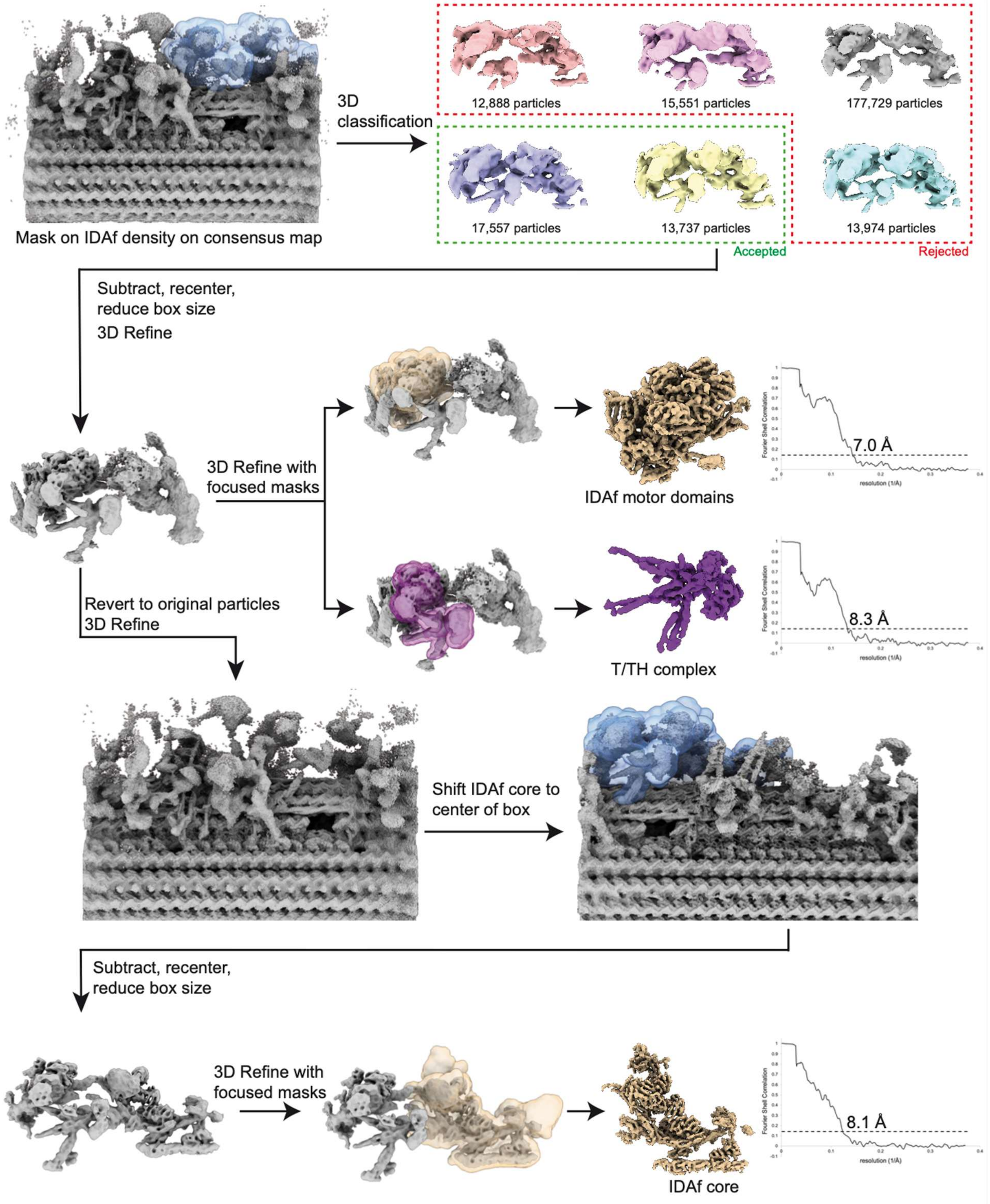

**Supplementary Figure 3 | Processing strategy for inner dynein arm *f* (IDA*f*).**
A total of 281,984 "96-nm half-2 particles" were subjected to 3D classification using a mask applied to diffuse density representing IDA*f*. Initial classification produced two well-defined classes that were combined and refined after subtraction. For final refinements, masks were created for the IDA*f* motor domains and the tetherhead complex, and local refinements were performed. To improve the IDA*f* core, the IDA*f* consensus particles were reverted to their original box size and centered on the core region for a consensus refinement. These recentered particles were then subtracted, followed by masked local refinement on the IDA*f* core.

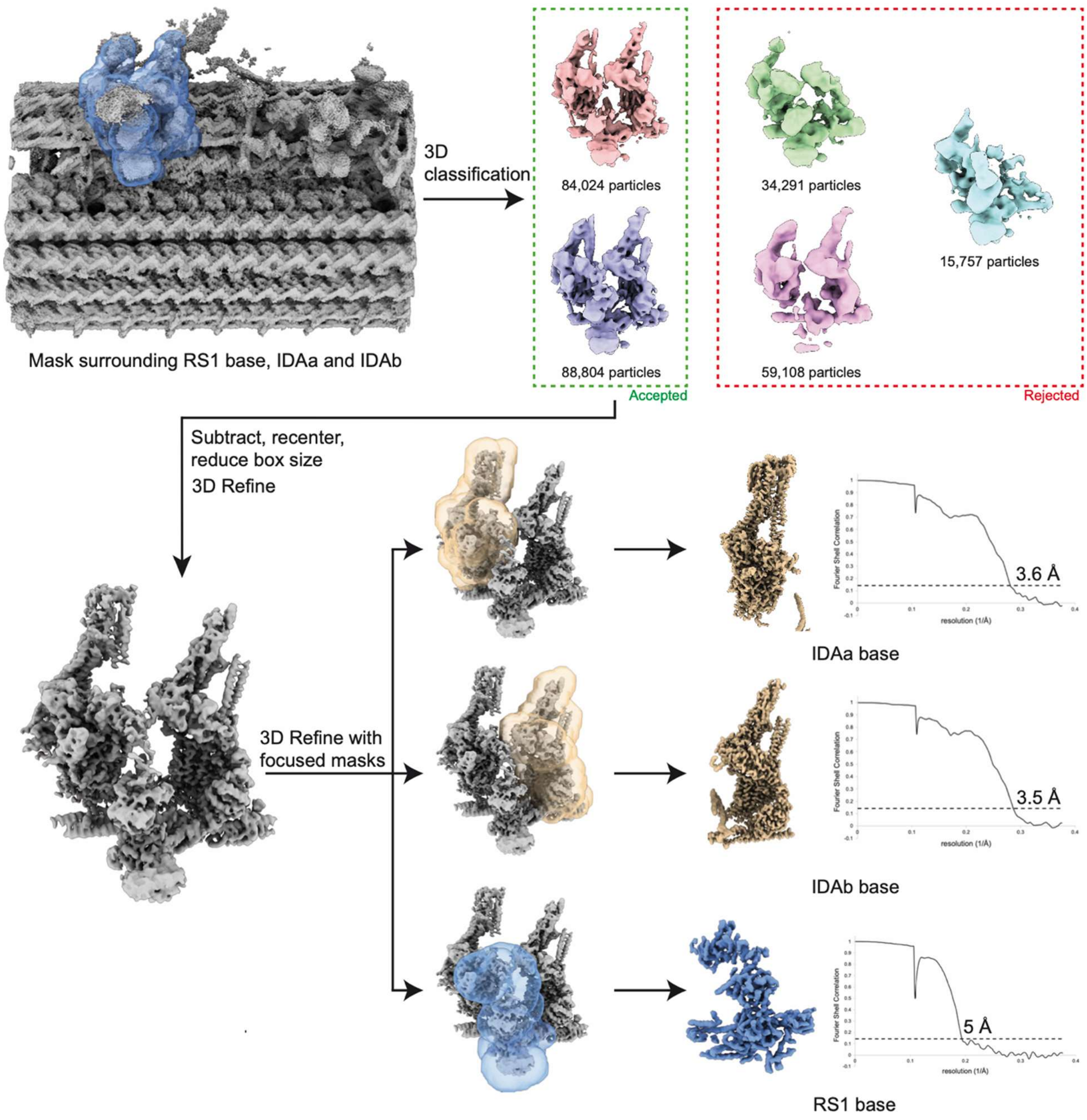

**Supplementary Figure 4 | Processing strategy for inner dynein arm (IDA) *a* and *b*.**
A total of 251,436 “96-nm half-1 particles” were subjected to 3D classification using a mask applied to diffuse density representing the base regions of radial spoke 1 (RS1), IDA*a*, and IDA*b*. Classification yield two classes containing well-resolved density that were combined and refined. To resolve individual regions, masks were created for the bases of IDA*a*, *b* and RS1 for use in final local refinements.

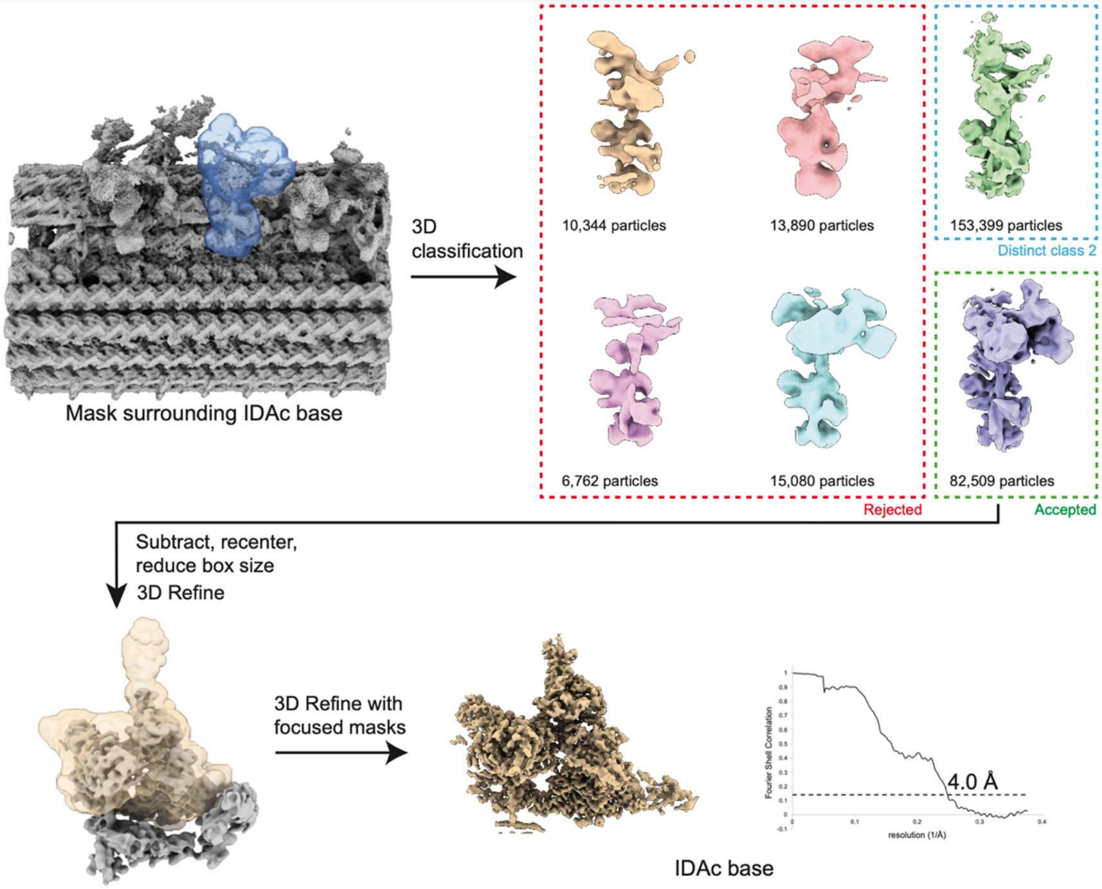

**Supplementary Figure 5 | Processing strategy for inner dynein arm *c* (IDA*c*).**
A shape mask was applied over diffuse density between RS1 and RS2. 3D classification revealed two well-resolved classes, one with 153,399 particles representing no IDAc and one with 82,509 particles representing IDA*c*. The class with IDA*c* was selected for re-refinement following signal subtraction, box-size reduction and recentering.

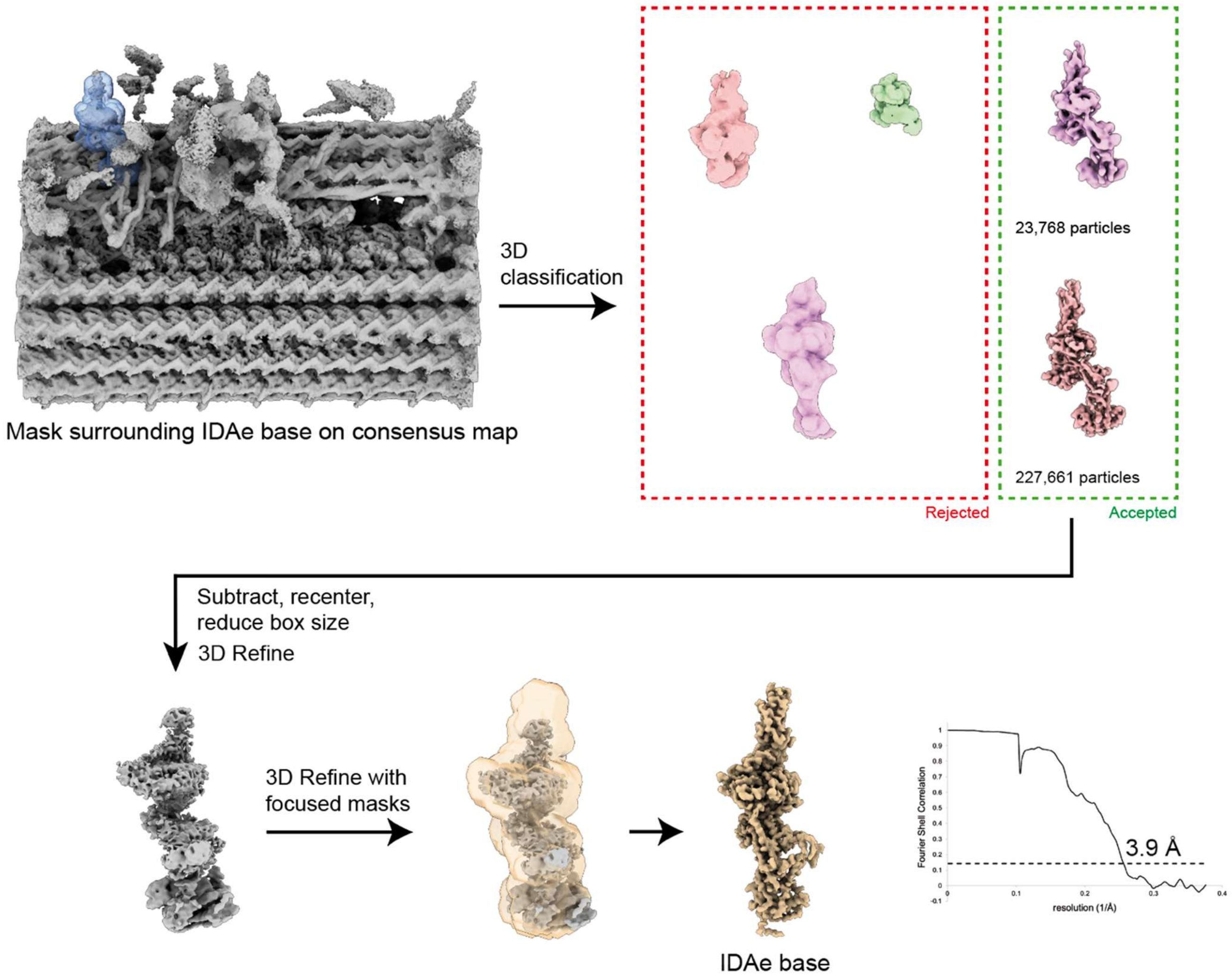

**Supplementary Figure 6 | Processing strategy for inner dynein arm *e* (IDA*e*).**
A shape mask was applied over diffuse density adjacent to the N-DRC. 3D classification revealed two well-resolved classes, containing a total of 251,429 particles. Particles in these classes were combined for re-refinement following signal subtraction, box-size reduction and recentering.

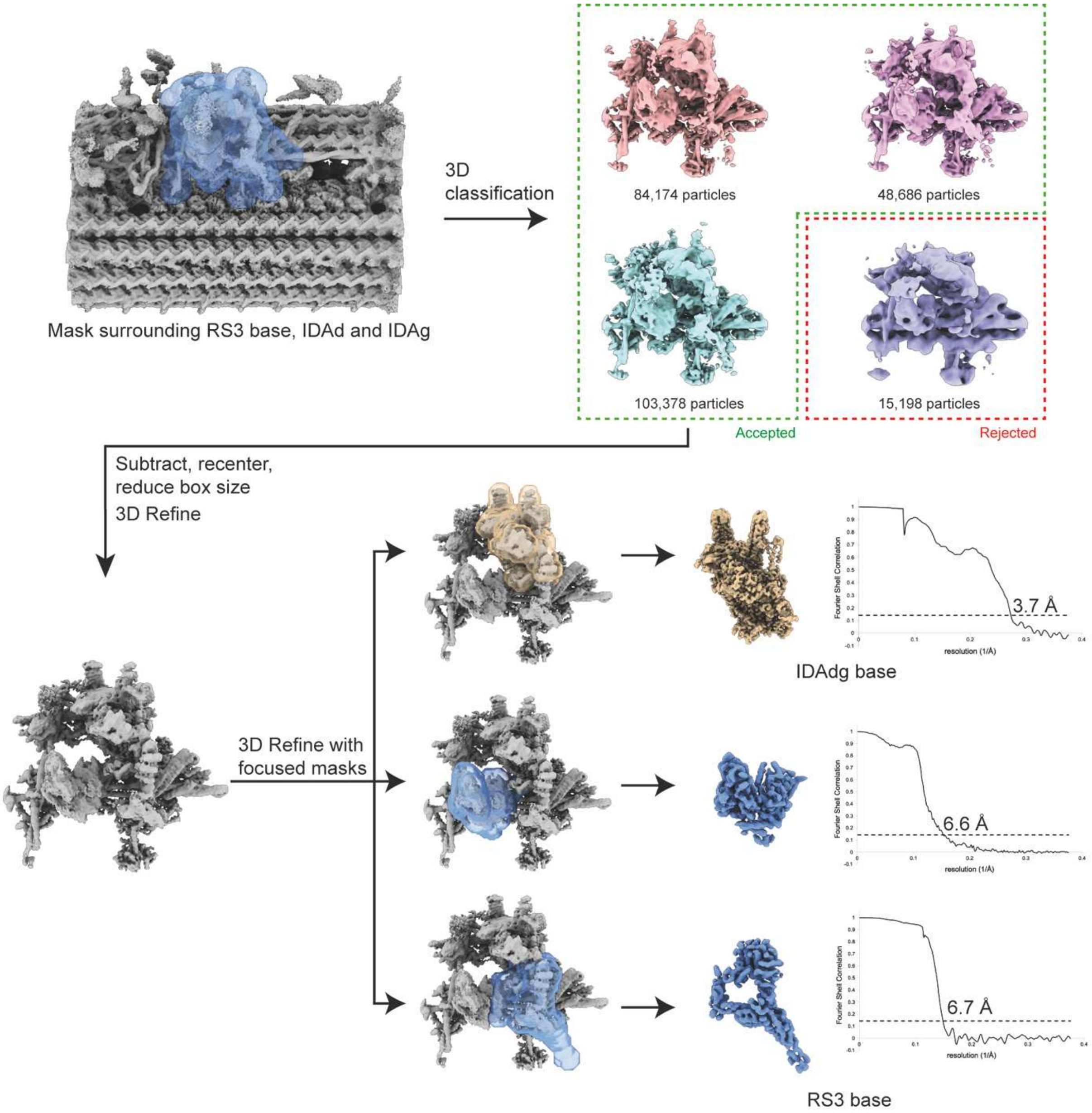

**Supplementary Figure 7 | Processing strategy for the bases of inner dynein arms *g, d*, and radial spoke 3 (RS3).**

A total of 281,984 “96-nm half-2 particles” were subjected to 3D classification using a mask applied to diffuse density representing the regions for the bases of radial spoke 3, IDA*g*, and *d*. Classification produced three classes containing well-resolved density that were combined and refined after subtraction. To resolve individual regions, masks were created for the base of IDA*g* and *d* and RS3 for use in final local refinements.

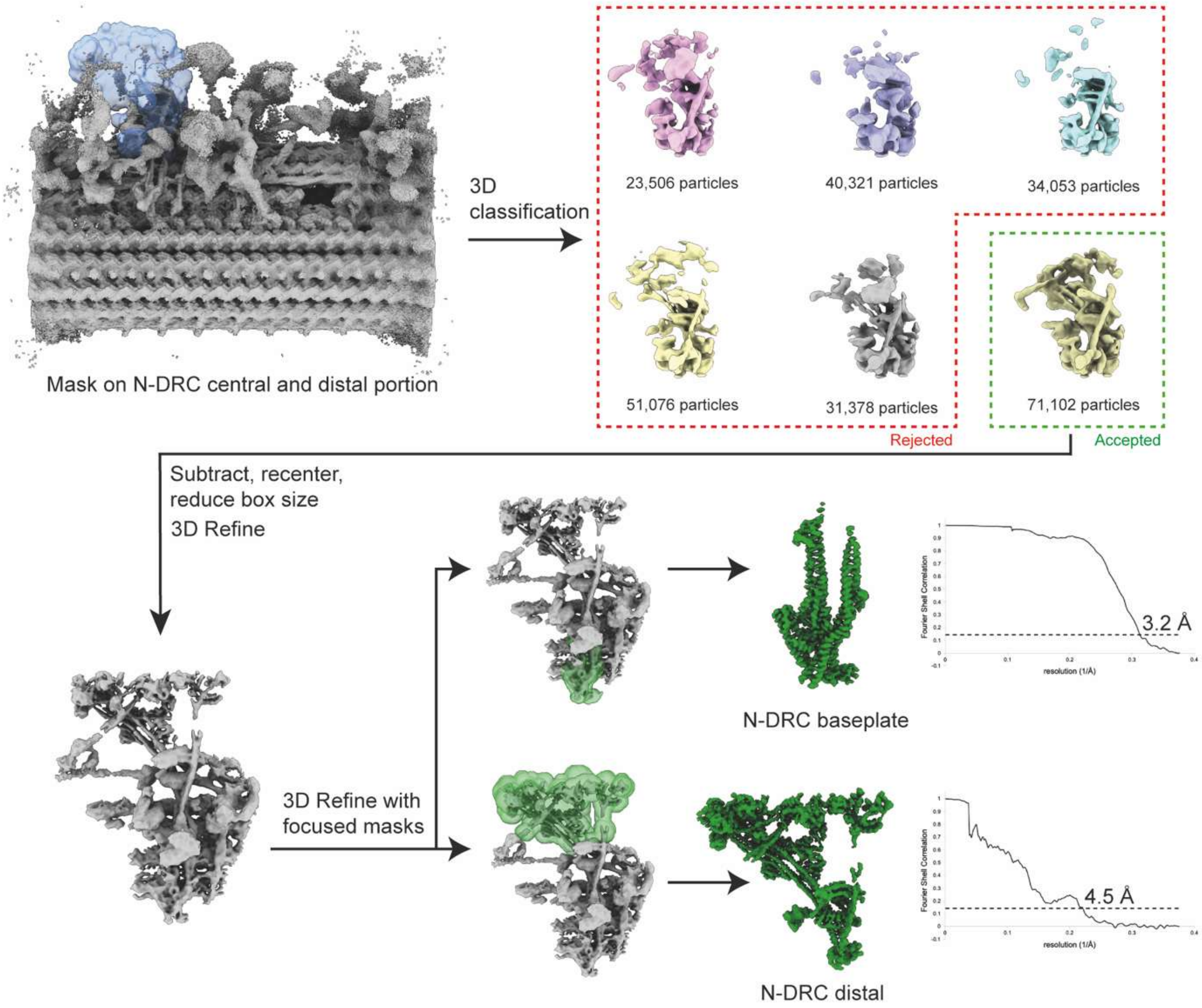

**Supplementary Figure 8 | Processing strategy for the nexin-dynein regulatory complex (N-DRC).**
A total of 281,984 “96-nm half-2 particles” were subjected to 3D classification using a mask applied to diffuse density representing the N-DRC. Classification yielded one class containing well-resolved density that was refined after subtraction to produce a consensus N-DRC refinement. To improve the quality and resolution of each region of the N-DRC, two masked were created for the N-DRC baseplate and N-DRC distal region. These were used in masked local refinements to produce the final reconstructions.

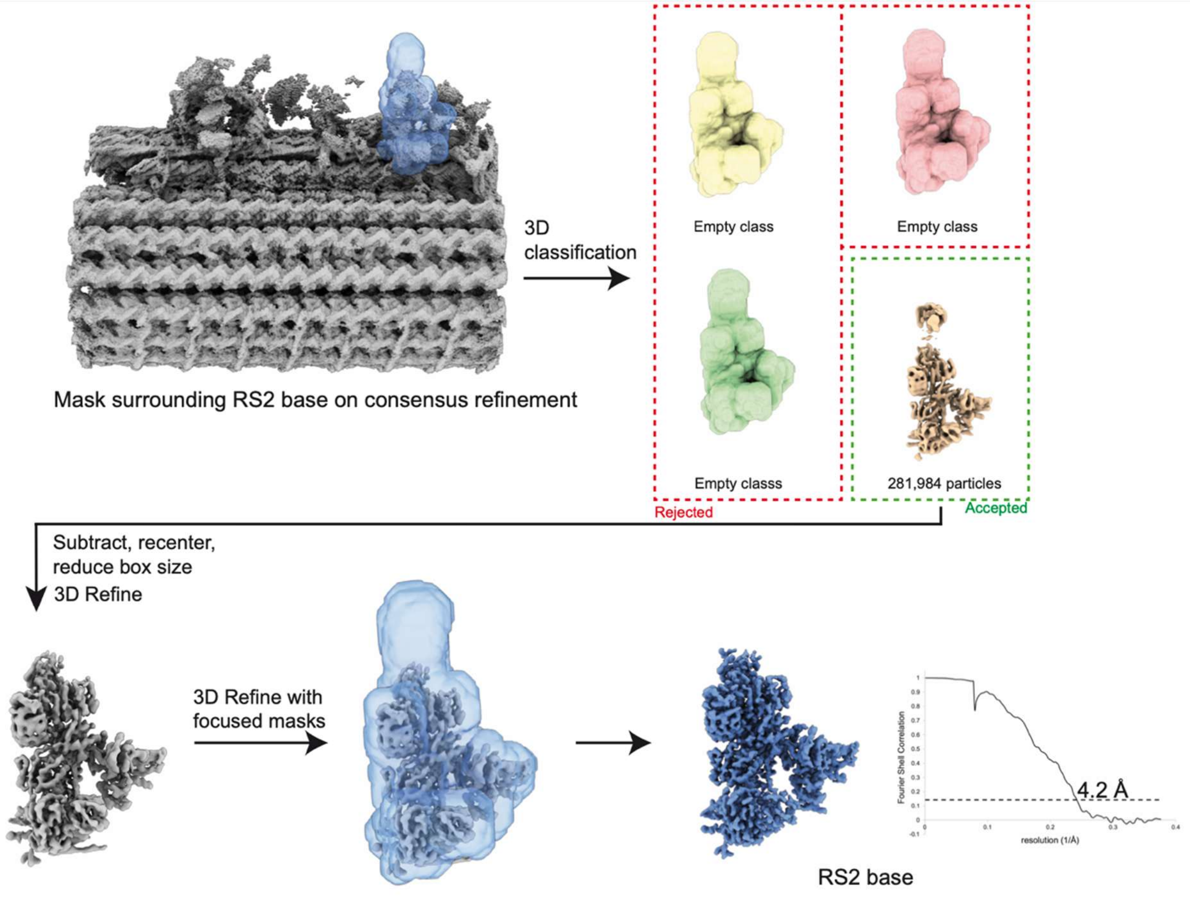

**Supplementary Figure 9 | Processing strategy for the base of radial spoke 2 (RS2).**
A total of 251,436 "96-nm half-1 particles" were subjected to 3D classification using a mask applied to diffuse density representing the regions for the bases of RS2. Classification produced one class with well-resolved density that was used for refinements after subtraction and recentering. A follow-up masked local refinement was performed to produce the final reconstruction for the RS2 base.

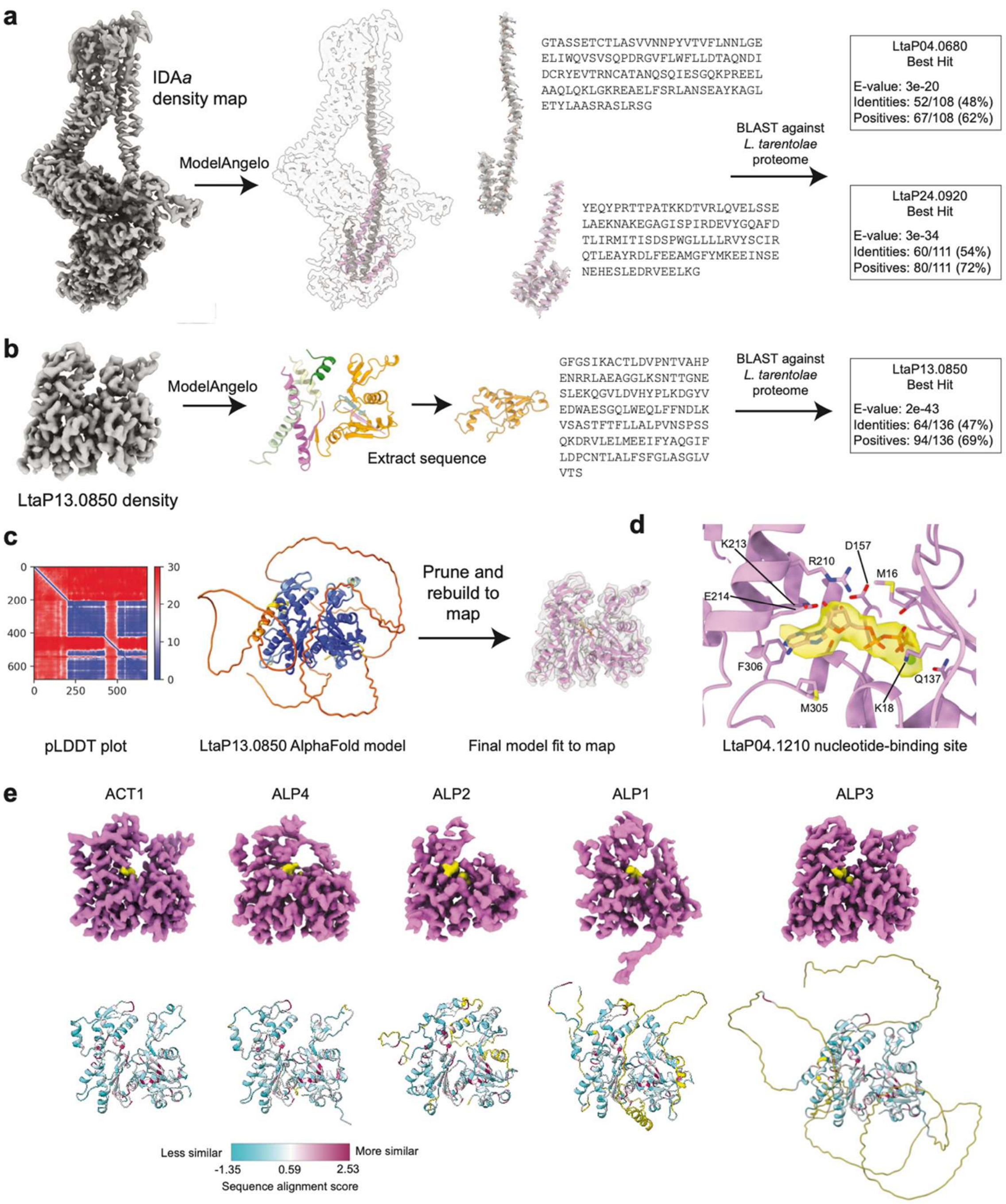

**Supplementary Figure 10 | Identifying DNALI1-like proteins (DLPs) and actin-like proteins (ALPs).**
**a**, DLPs within IDA*a* were identified using ModelAngelo. The two longest chains from the output were extracted and their sequences were compared against the *L. tarentolae* proteome using BLASTp. One hit with an outlier E-value was identified for each chain, corresponding to two DLP paralogs. **b**, ALPs were identified using ModelAngelo using the same approach as described in panel a. **c**, AlphaFold models were generated using the sequences identified for each ALP. The subsequent models were pruned to remove regions outside of the cryo-EM density and rebuilt to fit to the map. **d**, Cryo-EM density supporting an ATP-$Mg^{2+}$ bound in the ALP nucleotide-binding site. **e**, Top row: cryo-EM density maps for each ALP associated with single-headed IDAs. Bottom row: AlphaFold model for each ALP coloured by per-residue sequence conservation metrics estimated from a multiple sequence alignment using the AL2CO program in ChimeraX[16]. Divergent loops and terminal regions were not resolved in the cryo-EM density.

**Supplementary Tables 1 to 4**

**Table S1 | Proteins of the *Leishmania* axonemal 96 nm repeat**.
Table summarising the proteins associated with the 96-nm repeat of the axonemal doublet microtubule, grouped by structural complex. The table includes proteins identified in this study and in previous structures of trypanosomatid DMTs[17,18]. Note, only the centrin-like protein associated with IDA*g* (Centrin-5; LtaP32.0710) can be confidently assigned based on the density. Centrin proteins associated with IDA*c* and IDA*e* have been incorporated into the atomic model as LtaP07.0720 and LtaP36.6270. Provided as an Excel document.

**Table S2 | Dynein heavy chain identities.**
Gene identifiers (ID) for the dynein heavy chains from the three species analysed in this paper. *T. brucei* does not have a homolog of the *Leishmania* IDA*c* HC.

| **Dynein heavy chain** | ***L. tarentolae* gene ID** | ***L. mexicana* gene ID** | ***T. brucei* gene ID** |
|---|---|---|---|
| ODA-HCα | LtaP25.1030 | LmxM.25.0980 | Tb927.3.930 |
| ODA-HCβ | LtaP13.1510 | LmxM.13.1650 | Tb927.11.3250 |
| IDA*f*-HCα | LtaP34.3820 | LmxM.33.3880 | Tb927.4.870 |
| IDA*f*-HCβ | LtaP23.1600 | LmxM.23.1310 | Tb927.8.3250 |
| IDA*a* HC | LtaP27.2640 | LmxM.27.2590 | Tb927.2.5270 |
| IDA*b* HC | LtaPh_3609500 | LmxM.36.0950 | Tb927.10.5350 |
| IDA*c* HC | LtaP14.1050 | LmxM.14.1060 | |
| IDA*e* HC | LtaP26.0950 | LmxM.26.1020 | Tb927.7.920 |
| IDA*d* HC | LtaP28.0630 | LmxM.28.0610 | Tb927.11.8160 |
| IDA*g* HC | LtaP28.2900 | LmxM.28.2880 | Tb927.11.11220 |

**Table S3 | Swimming speed and directionality measurements.**
Swimming speed and directionality metrics for each analysed *L. mexicana* deletion strain. Deletion mutants for most dynein heavy chains, the ∆IC2 (LmxM.31.1060) and the ∆ODA-LC7A (LmxM.18.1010) were independently generated and analysed twice, with three measurements taken per experiment. LC4-like (LmxM.01.0620) was part of a pilot screen where only two speed measurements were recorded. Incomplete deletion denotes strains where the target open reading frame remained detectable by PCR, indicating that at least one percent of the population retained a copy of the gene. The columns “Growth”, “Motility”, “Curling”, “Shape of cells” and “Length of flagella” report qualitative observations of cells viewed in culture. Provided as an Excel document.

**Table S4 | Quantitative proteomics data of the 96 nm axonemal repeat proteins from *L. mexicana*.**
Protein abundances were determined using Top3 label-free quantification. The table presents raw and imputed Top3 intensities for individual axonemal proteins, along with calculated log2 fold changes relative to parental strains and Benjamini-Hochberg adjusted p-values for statistical significance. Provided as an Excel document.

### Supplementary References

1. Jamali, K. *et al.* Automated model building and protein identification in cryo-EM maps. *Nature* **628**, 450–457 (2024).

2. Walton, T. *et al.* Axonemal structures reveal mechanoregulatory and disease mechanisms. *Nature* **618**, 625–633 (2023).

3. Vagin, A. & Teplyakov, A. Molecular replacement with MOLREP. *Acta Crystallogr. Sect. D: Biol. Crystallogr.* **66**, 22–25 (2010).

4. Long, F., Vagin, A. A., Young, P. & Murshudov, G. N. BALBES: a molecular-replacement pipeline. *Acta Crystallogr. D Biol. Crystallogr.* **64**, 125–132 (2008).

5. Brown, A. *et al.* Tools for macromolecular model building and refinement into electron cryo-microscopy reconstructions. *Acta Crystallogr. Sect. D, Biol. Crystallogr.* **71**, 136–53 (2015).

6. Balasubramaniam, K., He, T., Chen, H., Lin, Z. & He, C. Y. Cytoplasmic preassembly of the flagellar outer dynein arm complex in Trypanosoma brucei. *Mol. Biol. Cell* **35**, br16 (2024).

7. Leung, M. R. *et al.* Structural diversity of axonemes across mammalian motile cilia. *Nature* **637**, 1170–1177 (2025).

8. Abramson, J. *et al.* Accurate structure prediction of biomolecular interactions with AlphaFold 3. *Nature* **630**, 493–500 (2024).

9. Beneke, T., Banecki, K., Fochler, S. & Gluenz, E. LAX28 is required for stable assembly of the inner dynein arm f/I1 and tether/tether head complex in Leishmania flagella. *J. cell Sci.* **133**, jcs239855 (2020).

10. Braga-Lagache, S. *et al.* Robust Label-free, Quantitative Profiling of Circulating Plasma Microparticle (MP) Associated Proteins*. *Mol. Cell. Proteom.* **15**, 3640–3652 (2016).

11. Wagner, T. M. *et al.* Extracellular vesicles of minimalistic Mollicutes as mediators of immune modulation and horizontal gene transfer. *Commun. Biol.* **8**, 674 (2025).

12. Pham, T. V., Henneman, A. A. & Jimenez, C. R. iq: an R package to estimate relative protein abundances from ion quantification in DIA-MS-based proteomics. *Bioinformatics* **36**, 2611–2613 (2020).

13. Silva, J. C., Gorenstein, M. V., Li, G.-Z., Vissers, J. P. C. & Geromanos, S. J. Absolute quantification of proteins by LCMSE: a virtue of parallel MS acquisition. *Mol. Cell. Proteom. : MCP* **5**, 144–56 (2005).

14. Huber, W., Heydebreck, A. von, Sültmann, H., Poustka, A. & Vingron, M. Variance stabilization applied to microarray data calibration and to the quantification of differential expression. *Bioinformatics* **18**, S96–S104 (2002).

15. Silver, J. D., Ritchie, M. E. & Smyth, G. K. Microarray background correction: maximum likelihood estimation for the normal–exponential convolution. *Biostatistics* **10**, 352–363 (2009).

16. Pei, J. & Grishin, N. V. AL2CO: calculation of positional conservation in a protein sequence alignment. *Bioinformatics* **17**, 700–712 (2001).

17. Xia, X. *et al.* Trypanosome doublet microtubule structures reveal flagellum assembly and motility mechanisms. *Science* **387**, eadr3314 (2025).

18. Doran, M. H. *et al.* Evolutionary adaptations of doublet microtubules in trypanosomatid parasites. *Science* **387**, eadr5507 (2025).
